# Recycling of uridylated mRNAs in starfish embryos

**DOI:** 10.1101/2022.04.07.487435

**Authors:** Haruka Yamazaki, Megumi Furuichi, Mikoto Katagiri, Rei Kajitani, Takehiko Itoh, Kazuyoshi Chiba

## Abstract

In eukaryotes, mRNAs with long poly(A) tails are translationally active, whereas deadenylation of the tails decreases translation and uridylation of the short poly(A) tails causes the mRNA to be degraded. In this study, we confirmed that maternal *cyclin B* mRNAs with long poly(A) tails in blastula embryos of invertebrate starfish were deadenylated and uridylated, followed by decay. In starfish oocytes, however, *cyclin B* mRNAs with uridylated short poly(A) tails are stable. They are polyadenylated and translationally active immediately following hormonal stimulation for resumption of meiosis. Similarly, maternal ribosomal protein mRNAs, *Rps29* and *Rpl27a*, which become uridylated following deadenylation upon hormonal stimulation, remain stable even after fertilisation and early development. At the morula stage, the uridylated maternal ribosomal protein mRNAs are modified to yield non-canonical poly (A) tails rich in U and G residues in the 5’ region and in A residues at the 3’ end, rendering them translationally active. These results indicate that the fates of uridylated mRNAs in starfish are decay and/or recycling.

## Introduction

In eukaryotes, most mRNAs carry a non-templated poly(A) tail at the 3’ end, modifications of which play important roles in regulating mRNA stability, transport, and translation. Long poly(A) tails can interact with poly(A) binding protein (PABP), which protects mRNAs from degradation and initiates translation by interacting with the 5’ cap-bound translation initiation factor 4F complex (eIF4F) (1). In humans, shortening of the poly(A) tail through the action of poly(A) nucleases such as the CCR4–NOT complex helps to remove PABP from the mRNA and subsequently to add uridine residues to the 3’ end by terminal uridylyl transferase 4 (TUT4) and TUT7 (2). The oligo(U) tails then recruit the LSM1–7 complex, degrading the mRNAs by the 5’ –3’ exonuclease XRN1 (2, 3). Uridylated mRNAs are also recognised and degraded by the 3’ –5’ exonuclease Dis3L2 (2, 4, 5). Similarly, in yeast *Schizosaccharomyces pombe*, a Dis3L2 homologue degrades uridylated RNA substrates (6). Therefore, deadenylation and uridylation of mRNA tails constitute a conserved system of RNA degradation in eukaryotes (7).

To accumulate the many ribosomes required for early embryonic development, ribosomal protein mRNAs that carry long poly(A) tails are translated within growing animal oocytes arrested at the prophase of meiosis I (Pro-I) (8). Following hormonal stimulation to resume meiosis, the ribosomal protein mRNAs are deadenylated to become translationally inactive, as observed in *Xenopus* oocytes (9). However, deadenylation of the mRNAs upon hormonal stimulation does not induce decapping or degradation during meiotic division; rather, the deadenylated mRNAs are stable until the blastula stage (9– 13), suggesting that additional steps may exist between deadenylation (or possibly uridylation) and decay of these mRNAs. In addition, the mechanism by which stability is conferred on transcripts with short poly(A) tails following deadenylation remains to be determined (14).

Poly(A) tails of maternal mRNAs in animal oocytes, such as *cyclin B*, are short, whereas they are stably stored. In *Xenopus* oocytes, the poly(A) lengths of *cyclin B1* mRNAs are controlled by cytoplasmic polyadenylation elements (CPEs) in their 3’ UTRs and the CPE binding protein (CPEB), recruiting poly(A) polymerase (Gld2) and deadenylating enzyme (PARN) (15–18). The poly(A) tails are kept short by PARN and Gld2 in dynamic equilibrium until hormonal stimulation (19). Upon meiosis resumption, CPEB phosphorylation occurs, which makes PARN release, causing poly(A) elongation and translational activation. Previously, we found that uridylation of *cyclin B* mRNA, which has a short poly(A) tail, does not cause RNA degradation in starfish oocytes arrested at Pro-I (20). After hormonal stimulation with 1-methyladenine (1-MA), which activates the Cdk1–cyclin B complex through the starfish serum- and glucocorticoid-regulated kinase (SGK)-dependent pathway (21, 22), some uridine residues of *cyclin B* mRNA were trimmed and the 3’ ends were subsequently polyadenylated (20). These results suggest that uridylation has a function other than the degradation of mRNA in starfish oocytes and that the mechanism of shortening poly(A) length differs from that of *Xenopus* oocytes. Moreover, it remains to be determined whether *cyclin B* mRNA degradation occurs following the uridylation concomitant with the maternal to zygotic transition (MZT) of starfish zygotes.

In this study, we investigated whether *cyclin B* mRNA was uridylated or degraded during starfish MZT. Furthermore, we studied the fate of ribosomal protein mRNAs during development to determine whether uridylation has a function other than mRNA decay.

## Materials and methods

### Animal and oocyte preparation

Starfish (*Asterina pectinifera*) were collected from the Pacific coast of Japan during the breeding season and were maintained in laboratory aquaria with seawater at 14 °C. To remove follicle cells, oocytes released from isolated ovaries were washed three times with ice-cold Ca^2+^-free seawater (450 mM NaCl, 9 mM KCl, 48 mM MgSO_4_, 6 mM NaHCO_3_, 40 mM 3-[4-(2-Hydroxyethyl)-1-piperazinyl]propanesulfonic acid (EPPS), pH 8.0) and incubated in artificial seawater (450 mM NaCl, 9 mM KCl, 48 mM MgSO_4_, 6 mM NaHCO_3_, 40 mM EPPS, 9.2 mM CaCl_2_, pH 8.0) or Jamarin artificial sea water (JAMARIN-U, Jamarin Laboratory, Osaka, Japan) at 20 ° C. Meiosis resumption of starfish oocytes was induced using 1 µM 1-MA. After verifying the breakdown of the germinal vesicle, “dry” sperm obtained from male starfish was added at a final dilution of 1/100,000, after which 1-MA and sperm were washed out. All experiments were performed at 20 °C, unless otherwise stated. Total RNA from oocytes was extracted using the RNeasy mini kit (QIAGEN, Hilden, The Netherlands). RNA was quantified using a Qubit 3.0 fluorometer (Thermo Fisher Scientific, Waltham, MA, USA).

### RT-PCR with 3’ adaptor ligation and tail sequence

Synthesis of cDNA with biotinylated 3’ adaptor ligation was performed using a Small RNA Cloning Kit (Takara Bio Inc., Kusatsu, Japan) as previously reported (20, 23). In some experiments, the thermostable group II intron reverse transcriptase (TGIRT) Template-Switching RNA-Seq Kit (InGex, Olivette, MO, USA) was used for reverse transcription (Fig. S1 B left panel). PCR was performed using gene-specific primers and a 3’ adaptor primer (primer sets are listed in Table S1). PCR products were purified by agarose gel electrophoresis and extracted from the gel using the Wizard SV Gel and PCR Clean-Up System (Promega, Madison, WI, USA). Purified PCR products were cloned into a pCR2.1-TOPO vector (Invitrogen, Carlsbad, CA, USA) and the insert sequences were determined by Sanger sequencing using Applied Biosystems™ 3130 DNA Analyzers (Applied Biosystems, Waltham, MA, USA).

### 5’-rapid amplification of cDNA

*Rps29* 5’UTR and coding region cDNAs were synthesised using SMART™ RACE cDNA Amplification Kit (Clontech Laboratories Inc. USA). PCR was performed using gene-specific primers (40S 5’ RACE R, Table S1) and a 5’ primer (SMART oligo) provided by the manufacturer. PCR products were purified and cloned into a pCR2.1-TOPO vector as described in the “RT-PCR with 3’ adaptor ligation and tail sequence” section.

### DNA cloning and plasmids for in vitro RNA synthesis

#### psfcycB_A25

To synthesise an RNA encompassing the 3’ UTR of starfish *cyclin B* carrying a unique sequence tag and a long poly(A) tail without uridines, we modified the pcDNA used in our previous study (20). A 25 nt poly(A) sequence was inserted after the 3’ UTR using the In-Fusion (Takara Bio) method.

#### psfRps29_WT

To create the pcDNA sfRps29 3’ UTR, the 3’ UTR of starfish *Rps29* cDNA was amplified by PCR using an sfRps29_F2 primer and PCR-RȦRT primer, and the product was TA-cloned into the vector pCR2.1-TOPO (Invitrogen, Carlsbad, CA, USA) as described in the “RT-PCR with 3’ adaptor ligation and tail sequence” section, followed another PCR. The starfish *Rps29* 3’ UTR cDNA was amplified using primers, psfRps29_WT_vector_F containing an exogenous sequence tag and psfRps29_WT_vector_R (fragment 1). Fragment 2 containing the 3’ region of the 3’ UTR of starfish *Rps29* cDNA, 20 bases of poly(A), and a BsmBI restriction endonuclease site was produced using PCR with psfRps29_WT_insert_template, and primers psfRps29_WT_insert_F and psfRps29_WT_insert_R. Fragment 3 was pBluescript KS (-) (Stratagene, California USA), which was digested using Xba. These three fragments were ligated using In-fusion (Takara Bio Inc, Shiga, Japan), and the resultant plasmid was named psfRps29_WT.

#### psfRps29_PAS

The Delta PAS sfRps29 vector was generated by PCR using PrimeSTAR MAX Polymerase (Takara Bio) with primers containing the point mutation (forward: 5’ - TCAGAAAGAAAATGACCAGATCTGCT-3’, and reverse: 5’ - TCATTTTCTTTCTGAACTCAATACAC-3’) and template psfRps29_WT. Point mutation sites in primers were underlined.

#### psfSGK vector

The psfSGK vector included the T7 promoter, the *A. pectinifera cyclin B* Kozak sequence (59-TACAAT-39), sfSGK (T479E), and a C-terminal 3× FLAG tag in the pSP64-S–based vector (21, 24, 25). To alter the 3’ terminal sequence and insert the BbsI site, we generated double-stranded DNA from single-stranded oligo DNAs via the 5’ –3’ polymerase activity of PCR polymerase, and inserted the product into the psfSGK vector using the In-Fusion method. The primers and oligo DNAs are listed in Table S2.

#### psf Rps29 UTR luciferase vector

*Rps29* 5’ UTR and coding region cDNAs in the pCR2.1-TOPO vector was amplified via PCR using primers sfRps29-GST_1F and sfRps29-GST_1R. The GST was amplified via PCR using primers sfRps29-GST_2F and sfRps29-GST_2R. The psfRps29 WT vector, including the 3’ UTR of *Rps29* mRNA, was amplified via PCR using primers sfRps29-GST_3F and sfRps29-GST_3R. They were ligated using In-fusion (Takara Bio Inc, Shiga, Japan) and the resultant plasmid was named psfRps29-GST. Then, the GST region of psfRps29-GST was replaced with luciferase cDNA (pNL1.1[Nluc]Vector; Promega Madison, WI, USA). Luciferase cDNA was amplified via PCR using primers Nluc_FW2 and Nluc_RV2. The psfRps29-GST was amplified via PCR using primers vec_FW2 and vec_RV2. These PCR fragments were ligated using In-fusion and the resultant plasmid was named psf Rps29 UTR Luciferase, which was used to produce the reporter mRNA (Fig. 6. C upper panel).

### Microinjection

Microinjection was performed as previously described (20, 26). Briefly, the oocytes were injected using a constricting pipet filled with in vitro synthesised RNA (10-100 ng/µL) (injection volume, 1–2% oocyte volume). Injected oocytes were incubated for the indicated periods; 100 oocytes were used for each experiment.

### In vitro transcription

In vitro transcription of starfish *cyclin B* (*sfcyclin B*) mRNA was performed as previously described (20) using the mMESSAGE mMACHINE T7 kit (Invitrogen) with the original NTP concentrations. Transcripts were subsequently polyadenylated using the Poly(A) Tailing Kit (Thermo Fisher Scientific) and purified using phenol/chloroform extraction and ethanol precipitation.

The DNA templates for in vitro transcription of sfRps29_WT mRNA and sfRps29_delta PAS mRNA were PCR-amplified from the pRps29_WT vector or pRps29_PAS vector, respectively, using a universal M13 forward primer (5’ - GTAAAACGACGGCCAGT-3’) and universal M13 reverse primer (5’ - CAGGAAACAGCTATGAC-3’). The PCR product was digested with BsmBI (Esp3I), and the desired fragment was purified by agarose gel electrophoresis using the Wizard SV Gel and PCR Clean-Up System followed by phenol/chloroform extraction and ethanol precipitation, with the resultant DNA pellet subsequently dissolved in sterile water. Using this DNA as a template, in vitro transcription was performed using the mMESSAGE mMACHINE T7 kit with the original NTP concentrations. Transcripts were subsequently polyadenylated using the Poly(A) Tailing Kit and purified by phenol/chloroform extraction and ethanol precipitation.

The DNA templates for SGK were amplified using the psfSGK_RNA_F primer (5’ -GCCAGATGCTACACAATTAGGC-3’) and psfSGK_RNA_R primer (5’ - TTGTTTGCCGGATCAAGAGC-3’). PCR products were digested with BbsI (BpiI); polyadenylation using a Poly(A) Tailing Kit was performed when long poly(A) was required.

The psf Rps29 UTR Luciferase vector was digested with BsmBIand purified via agarose gel electrophoresis using the Wizard SV Gel followed by phenol/chloroform extraction, ethanol precipitation, in vitro transcription, and poly(A) elongation as mentioned above.

### Real-time qPCR

Total RNA was extracted using the RNeasy Micro Kit (QIAGEN) and 300 ng reverse transcribed using an iScript select cDNA synthesis kit (Bio-Rad, Berkeley, CA, USA) using random hexamers. The cDNAs were quantified with SYBR-Green assays (Applied Biosystems, Foster City, CA, USA) using the Applied Biosystems 7500 Real-Time PCR System; 18S rRNA was used for normalisation. The primers used for qRT-PCR are listed in Table S2.

### Translation assay

SGK mRNAs were injected into the oocytes. Oocytes incubated in Jamarin seawater for the indicated periods were recovered in 3 µL of seawater, added directly to 3 µL of 2× Laemmli sample buffer, and immediately frozen in liquid N_2_. After thawing and boiling for 5 min at 95 °C, the proteins were separated on 8% polyacrylamide gels and immunoblotted (primary antibody: anti-sfSGK-HM (1:1,000, in TBS-T) (21, 22); secondary antibody: HRP-conjugated anti-rabbit IgG (A6154, Sigma; 1:2,000, in TBS-T)). Proteins recognised by the antibodies were visualised using ECL prime (GE Healthcare, Chicago, IL, USA) and digital images were acquired with a LAS4000 mini imager (FUJIFILM Wako Pure Chemical, Osaka, Japan).

### Luciferase assay

Eggs and embryos were lysed using Reporter Lysis Buffer (Promega, Madison, WI, USA). Luminescence was obtained using Nano-Glo®□ Luciferase Assay System (Promega, Madison, WI, USA) and quantified with Lumicounter 700 (MICROTEC CO., LTD, Funabashi, Japan).

### Illumina MiSeq sequencing for targeted TAIL-Seq

The multiplexed cDNA samples were sequenced using Illumina MiSeq (San Diego, CA, USA) (Table S3). Sequencing was performed to obtain paired-end reads (2 × 300 nt) according to the manufacturer’s protocol. The quality filter was turned off so as not to remove reads comprising poly(A), which often include low-quality bases. To prevent saturation of similar sequences, DNA samples from other organisms were mixed with the starfish-derived cDNAs to decrease the ratio of the cDNA samples to 22%. Note that reads from these organisms had distinct sequences from the target cDNA of the starfish and therefore could be separated via different barcodes (indices) and excluded from downstream analysis.

### Correction of mRNA (cDNA) sequences for mapping of targeted TAIL-Seq reads

To increase mapped reads and prevent misdetection of 3’ tail modifications, mRNA (cDNA) sequences used as reference sequences for read mapping were modified into those of individual samples (Supplemental Figure S1A1). The target genes were *Rps29* (40S ribosomal protein) and *L27a* (60S ribosomal protein). The procedure for each sample was as follows.

1. To prevent misalignments of reads, a poly(A) sequence of 50 nt length was added to the 3’ end of mRNA in silico.
2. The reads of the GV sample from oocytes before hormonal stimulation were randomly downsampled to a coverage depth of 1000×.
3. The downsampled reads were mapped to the mRNA sequence using BWA-MEM (27) (version 0.7.12-r1044) with default parameters.
4. The mapped reads were piled up using SAMtools (28) (version 1.3.1). The commands were ‘sort’, ‘index’, and ‘mpileup’ with the parameters, and the output was the text file of the pileup result.
5. For each position in the mRNA sequence, a base (A, T (U), G, or C) was changed to the majority position using the pileup result and the in-house program.
6. The poly(A) sequence added to (**1**) was removed.

### Extraction of valid 3’ reads of targeted TAIL-Seq

For each sample, the procedures were as follows (the processes without tool names were performed using in-house programs) (Supplemental Figure S1A2).

1. The 3’ adopters in reads were searched using BLASTN (29) (version 2.6.0) with the option of ‘-task blastn-short -word_size 6 -window_size 0 -dust no’ to handle short sequences. Here, the query and database inputs were reads and adopters, respectively (‘-query reads.fa -db adopter’).
2. Reads in which 3’ adopters were aligned with identity ≥90%, alignment coverage = 100%, and alignment positions were the heads of reads were extracted. Read pairs without the 3’ adopter were discarded. For each extracted read pair, the reads with and without the 3’ adopter were treated as 3’ and 5’ reads, respectively.
3. The 5’ reads were aligned to the targeted mRNA using BLASTN with the options of ‘task blastn -word_size 10 -window_size 0 -dust no’ read pairs in which 5’ reads were aligned with identity ≥90% and alignment length ≥200 nt were extracted, and the others were discarded to exclude contamination.

### Analysis of poly(A) and 3’ modifications in targeted TAIL-Seq reads

The procedures were based on the methods introduced in the initial TAIL-Seq study (30) with some modifications (Supplemental Figure S1A3, A4). Reads were treated as FASTQ files, not applying the previous methods utilising raw signal information, to analyse non-canonical poly(A) sequences. Poly (A) was sequenced as poly(T) because 3’ reads corresponded to the reverse complements of mRNA sequences. The following procedures targeted the reverse complements of 3’ reads to provide concise descriptions and were performed using in-house programs.

1. The 3’ reads of the extracted pairs (see ‘Extraction of 3’ reads of targeted mRNAs’) were aligned to the targeted mRNA using BLASTN with the options of ‘-task blastn - word_size 10 -window_size 0 -dust no’.
2. If an alignment with length ≥30 nt was detected in a 3’ read, the aligned region was treated as a 3’ UTR (not a 3’ tail region). To handle low-quality reads associated with poly(A), a sequence identity cutoff was not applied.
3. To detect poly(A) regions, a score was calculated based on the composition of bases as follows: A, +1; N, −1; T, G, or C, −2. If the score was higher, it was considered that the region was more likely to comprise poly(T). Although the concept of this score was introduced in a previous study (30), our calculation method differed from that originally published (A, +1; N, −2; T, G, or C, −10) to detect non-canonical poly(A) regions. For each 3’ read, the scores were calculated for all regions (substrings) and the region that satisfied the following conditions was determined as poly(A): score was maximum through the read, score >0, and a distance to the 3’ end of the read ≤15 nt.
4. Using the results of (**2**) and (**3**), regions in each 3’ read were classified into 3’ UTR, poly(A), 3’ end modification (a region between poly(A) and the 3’ end of a read), and other. Statistics, such as length, were calculated for each class. When calculating the length distributions of poly(A)s, the regions with lengths ≥40 nt were treated as the same category (‘≥40 nt’) because lengths of long poly(A)s tended to be overestimated due to systematic base-calling errors (30).
5. For each 3’ end modification, classification was based on the majority base (T, G, or C). If multiple bases occurred at the same time, the modification was classified as ‘≥2’.
6. To reduce the influence of sequencing errors, bases in 3’ reads with quality values <20 were masked, and compositions of bases (rates of A, T, G, and C) were calculated using unmasked reads.

### Analysis of TAIL-Seq data of *Xenopus laevis*

The TAIL-Seq data for *X. laevis* (African clawed frog) were generated in a previous study (31). The targeted sample set comprised the wild-type *X. laevis* early embryos (internal ID, hs27), which comprised embryos from the zygote (1 cell) to stage 12. The procedures were as follows (processes without tool names were performed using in-house programs).

1. The data set was generated and pre-processed in a previous study (31) using Tailseeker (version 3.1.7; https://github.com/hyeshik/tailseeker); the pre-processing steps included removal of adaptors and PCR duplicates. Intermediate FASTQ files of paired reads were obtained through personal communication with Dr. Hyeshik Chang (an author of the previous study).
2. 5’ reads were mapped to the *X. laevis* RNA sequence set (RefSeq accession, GCF_001663975.1; file name, GCF_001663975.1_Xenopus_laevis_v2_rna_from_genomic.fna) using BWA-MEM (27) (version 0.7.12-r1044) with default parameters.
3. The 5’ reads that were primarily mapped to the mRNA of rps29 (40S ribosomal protein; accession, NM_001171730.1) were detected and the corresponding 3’ reads extracted.
4. For the extracted 3’ reads, the same procedures were applied as for the starfish (see ‘Detection and analysis of 3’ ends of mRNA including poly(A)’).

In addition, we also attempted to analyse the TAIL-Seq data from *X. laevis* oocytes (internal ID, ms97); however, the number of reads that mapped to the *rps29* mRNA was too small to analyse.

## Results

### Decay of *cyclin B* mRNA occurs following deadenylation and uridylation at MZT

Deadenylation and uridylation of maternal mRNAs, including *cyclin B2* in *Xenopus* and zebrafish embryos, causes decay of mRNAs at MZT (31, 32). In starfish, the expression of many zygotic genes, including *wnt8* and *foxq2*, that constitute marker genes of MZT, is initiated at the blastula stage (33–35).

To determine whether maternal *cyclin B* mRNAs in starfish are degraded during meiosis resumption, fertilisation, or MZT, we used real-time qPCR (RT-qPCR) to monitor maternal *cyclin B* mRNA expression levels before fertilisation in Pro-I oocytes and in MI oocytes forming the first polar body (1PB), or after fertilisation in embryos at the morula, blastula, and gastrula stages. We found that maternal *cyclin B* mRNA levels were decreased in gastrula embryos (Figure 1A), suggesting that the decay of *cyclin B* mRNAs occurred at the MZT between the blastula and gastrula stages. In addition, these results confirmed our previous conclusion that *cyclin B* mRNAs in Pro-I oocytes do not decrease following meiosis resumption, although they carry uridylated short poly(A) tails (20).

**Figure 1.**
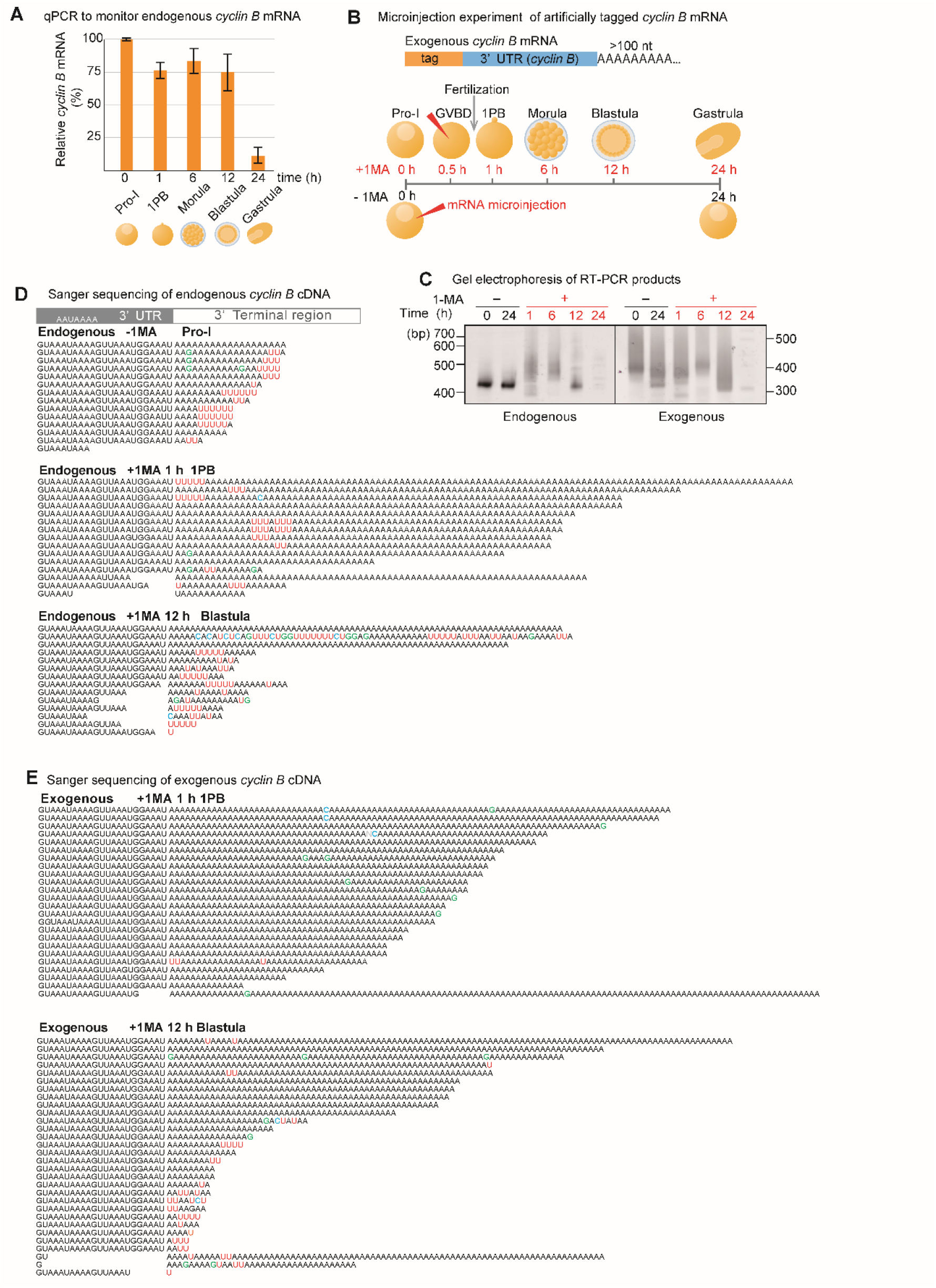
*Cyclin B* mRNA uridylation and degradation. A. Relative mRNA expression level of *cyclin B* in starfish oocytes and embryos. Total RNAs were purified at the indicated time and relative quantification of *cyclin B* mRNA expression was conducted using RT-qPCR (median ± SEM) (n = 3). B. Experimental scheme for microinjection of artificially tagged *cyclin B* mRNA. Tag-labelled *cyclin B* mRNA was synthesised and injected (red arrow heads) into oocytes at 0.5 h after 1-MA treatment (+1-MA) or into oocytes without 1-MA treatment (−1-MA). Oocytes (+1-MA) were inseminated before the first polar body formation and total RNAs were purified at the indicated time. C. Measurement of tail lengths of *cyclin B* mRNA. At the indicated time, the total RNAs were purified, and TGIRT template-switching reaction was performed (Supplemental Figure S1, B left panel). RT-PCR was conducted using the 3’ adaptor reverse primer and *cyclin B*-specific forward primer. The PCR products were then subjected to polyacrylamide gel electrophoresis and visualised using SYBR-Green I staining. Left and right panels show changes in the tail lengths of endogenous *cyclin B* mRNA and exogenously microinjected *cyclin B* mRNAs, respectively. D. Sanger sequencing results of the 3’ terminal region of cDNA from endogenous maternal *cyclin B* mRNA of oocytes at Pro-I without 1-MA treatment [Endogenous −1-MA], from stimulated oocytes following first polar body formation (1 h after 1-MA treatment) [Endogenous +1-MA 1 h], and from blastula embryos (12 h after 1-MA treatment) [Endogenous +1-MA 12 h]. E. Sequencing results of the 3’ terminal region of exogenous *cyclin B* mRNA from stimulated oocytes following first polar body formation (1 h after 1-MA treatment) [Exogenous +1-MA] and from blastula embryos (12 h after 1-MA treatment) [Exogenous −1-MA].

To monitor tail lengths and sequences of the endogenous maternal *cyclin B* mRNAs during meiosis resumption, fertilisation, and development, we produced cDNAs containing an adapter ligated to the 3’ ends using a template-switching reaction catalysed by the TGIRT-III enzyme (Supplemental Figure S1, B left panel). We then performed RT-PCR using the 3’ adaptor primer and a *cyclin B*-specific primer designed to hybridise 373 nt upstream of the polyadenylation site (sfcycB F primer; Table S1). Polyacrylamide gel electrophoresis of the RT-PCR products, including the 3’ adapter sequence, yielded a single 420 bp band from oocytes at Pro-I (Figure 1C, left panel, endogenous). This band was stably detected in unstimulated oocytes incubated in seawater for 24 h after isolation. In comparison, after hormonal stimulation, 450 to 550 bp mobility shifted bands were produced (Figure 1C, left panel: 1 and 6 h, endogenous). At the blastula stage after 12 h of hormonal stimulation, shorter bands appeared (Figure 1C left panel, 12 h, endogenous), followed by disappearance of the band in gastrula stage embryos (Figure 1C, left panel, 24 h, endogenous).

As previously reported (20), Sanger sequencing of the RT-PCR product showed that endogenous *cyclin B* mRNA in oocytes before hormonal stimulation carried uridylated short poly(A) tails (Figure 1E, endogenous −1MA), which were polyadenylated after hormonal stimulation (Figure 1D, endogenous +1-MA 1 h). These long poly(A) tails were stable until the morula stage (Figure 1C, left panel, 6 h, endogenous; Supplemental Figure S1D, Endogenous, +1-MA 6 h Morula). Furthermore, at the blastula stage, short poly(A) tails were uridylated (Figure 1D, endogenous +1-MA 12 h), followed by decay of the *cyclin B* mRNAs at the MZT between the blastula and gastrula stages (Figure 1A).

Because some uridine residues originating from oligo(U) tails at Pro-I oocytes were included in the long polyadenylated tails of the *cyclin B* mRNA following hormonal stimulation (20) (see also Figure 1D, endogenous +1-MA 1 h 1PB), they could be exposed at the 3’ end of the tails through trimming or partial deadenylation of the long poly(A) at the 12 h blastula stage. To distinguish these ‘old’ uridines exposed upon deadenylation at the blastula stage from possible ‘new’ uridylation during development, we synthesised an exogenous RNA encoding the 3’ UTR of starfish *cyclin B* carrying both a unique sequence tag and a long poly(A) tail without uridines (Figure 1B, upper panel). When we injected this RNA into the oocytes following hormonal stimulation, the lengths of RT-PCR products from this exogenous mRNA became shorter at the blastula stage and disappeared at the gastrula stage (Figure 1C, right panel), showing a pattern comparable to that of the endogenous *cyclin B* mRNA (Figure 1C, left panel). In addition, Sanger sequencing revealed that exogenous mRNAs were deadenylated and uridylated at the 12 h blastula stage (Figure 1E), followed by decay of the mRNA (Figure 1C, right panel). These results suggest that uridylation of *cyclin B* mRNA at the blastula stage in starfish embryos may function to induce degradation of the mRNAs, as previously demonstrated in vertebrate embryos (31) and that the 3’UTR of *cyclin B* is sufficient for deadenylation and uridylation of the mRNA.

### Ribosomal protein mRNAs are uridylated after 1-MA stimulation and fertilisation, followed by non-canonical poly(A) tail formation in blastula embryos

Because ribosomes are required for translation of maternal mRNAs such as *cyclin B* mRNA, mRNAs of ribosomal proteins carrying long poly(A) tails are actively translated in oocytes to produce ribosomes before hormonal stimulation. Following hormonal stimulation to resume meiosis, the deadenylation of ribosomal protein mRNAs leads to the cessation of ribosome production, as shown in *Xenopus* oocytes (8, 9), although the mRNA of the ribosomal protein L5 or L13 was still detected until stage 6 of the early blastula (36, 37). These results suggest that deadenylation of ribosomal protein mRNAs does not always cause the decay of mRNAs. Indeed, expression levels of starfish ribosomal protein mRNA (*Rps29*), evaluated using RT-qPCR, did not decrease in stimulated oocytes or in embryos at the morula and blastula stages (Figure 2A). Instead, mRNA levels increased in blastula embryos, which was likely due to the zygotic expression of *Rps29* mRNA, as previously shown in *Xenopus* embryos (37).

**Figure 2.**
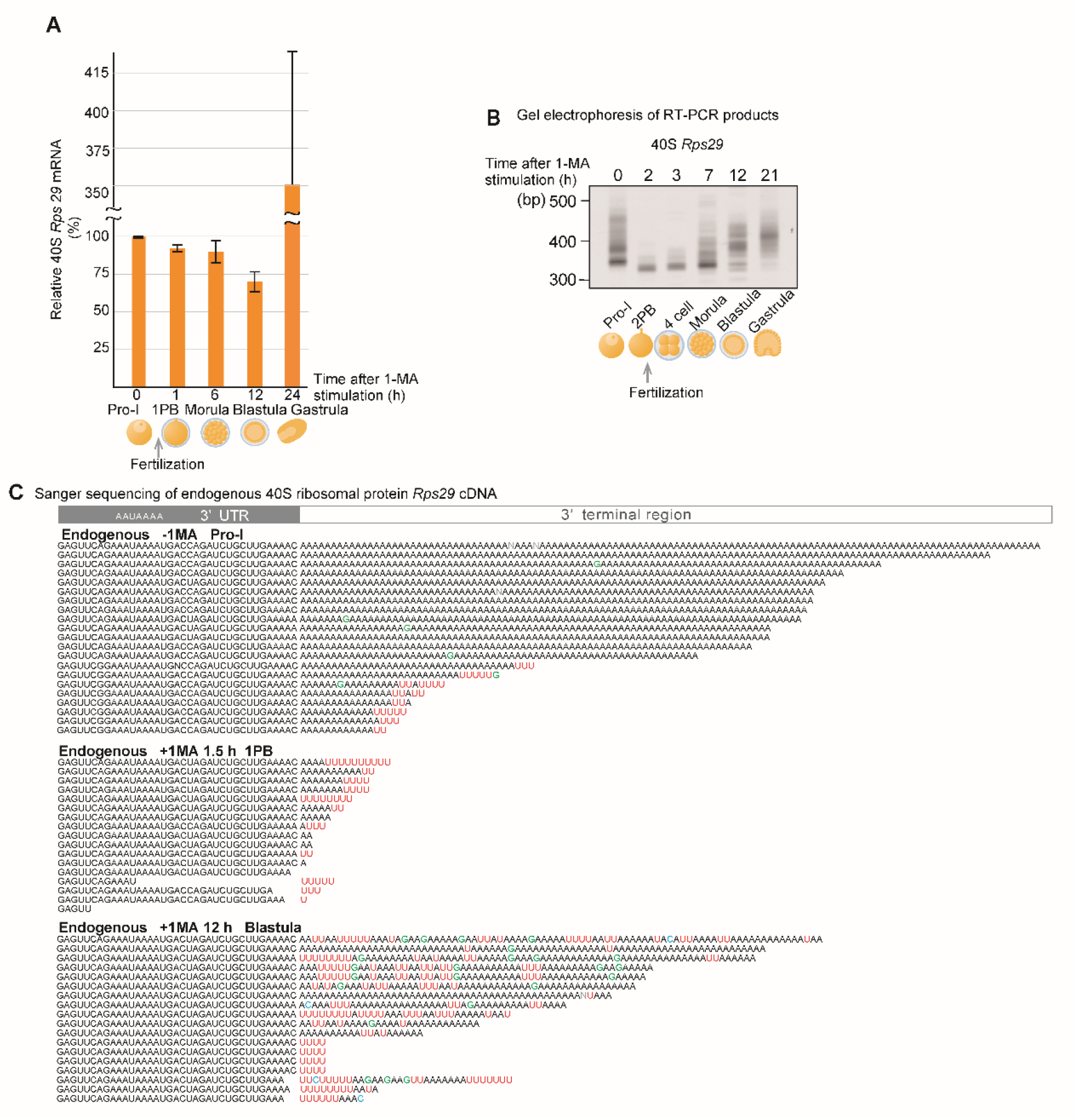
Deadenylation, uridylation, and non-canonical poly(A) elongation of the ribosomal protein *Rps29* mRNA. A. Relative levels of mRNA expression of ribosomal protein in starfish oocytes and embryos. Total RNAs were purified at the indicated time and relative quantification of ribosomal protein *Rps29* mRNA expression was performed using RT-qPCR (median ± SEM) (n = 3). B. Measurement of tail lengths of *Rps29* mRNA. Stimulated oocytes with 1-MA were inseminated following second polar body formation. At the indicated time before or after 1-MA stimulation, total RNAs were purified and adaptor ligation performed (Supplemental Figure S1, B right panel). RT-PCR was conducted using the 3’ adaptor reverse primer and *Rps29*-specific forward primer. The PCR products were then subjected to polyacrylamide gel electrophoresis and visualised using SYBR-Green I staining. Similar results are obtained using 3 animals. C. Sequencing results of the 3’ terminal region of cDNA of endogenous maternal *Rps29* mRNA from oocytes at Pro-I without 1-MA treatment [Endogenous −1-MA], stimulated oocytes after the first polar body formation (1.5 h after 1-MA treatment) [Endogenous +1-MA 1.5 h], and blastula embryos (12 h after 1-MA treatment) [Endogenous +1-MA 12 h].

To determine whether the 3’ ends of the ribosomal protein mRNAs of starfish oocytes or embryos were modified during the resumption of meiosis and early development, we purified total RNAs at the indicated times (Figure 2B) and then ligated a 3’ adaptor to the 3’ ends of the total RNAs (20) (Supplemental Figure S1, B right panel). When we amplified the 3’ end of the 40S ribosomal protein *Rps29* mRNA using a gene-specific primer and a 3’ adaptor primer, polyacrylamide gel electrophoresis of the RT-PCR products showed that the broad bands (320–470 bp) apparent in oocytes before hormonal stimulation (Figure 2B, 0 h after 1-MA stimulation) became shorter and sharper (310 bp) in oocytes following hormonal stimulation (Figure 2B, 2 h after 1-MA stimulation). This shortening of PCR product lengths may be due to deadenylation induced by 1-MA stimulation, as decreases in the poly(A) lengths of ribosomal proteins are well reported in *Xenopus* oocytes after meiotic resumption (9). Consistent with this, Sanger sequencing showed that more than half of *Rps29* mRNAs before hormonal stimulation contained long poly(A) tails (Figure 2C, −1-MA), whereas all mRNAs in stimulated oocytes carried short poly(A) or uridylated short poly(A) tails (Figure 2C, +1-MA, 1.5 h). Furthermore, even before hormonal stimulation, some *Rps29* mRNAs carried short uridylated poly(A) tails (Figure 2C, −1-MA), which were likely to become shorter after hormonal stimulation (Figure 2C, +1-MA, 1.5 h).

Notably, the lengths of the RT-PCR products became longer in morula and blastula embryos (Figure 2B), whereas the amount of mRNA did not increase at these stages (Figure 2A). Sanger sequencing of the PCR products revealed that the long poly(A) tails of mRNAs from blastula embryos included many U and some G nucleotides (Figure 2C, +1-MA, 12 h). We defined such tails as non-canonical poly(A) tails.

Similarly, the mRNA of 60S ribosomal protein L27a, *Rpl27a*, was deadenylated and uridylated after hormonal stimulation (Supplemental Figure S2, −1-MA; +1-MA, 1.5 h). In blastula embryos, the *Rpl27a* mRNAs carried non-canonical poly(A) tails including many GU nucleotides (Supplemental Figure S2, +1-MA, 12 h).

### Targeted TAIL-Seq of *Rps29* mRNA

Next, to obtain more detailed quantitative and qualitative information about the tail structures of *Rps29* mRNA, we modified the ‘TAIL-Seq’ method by Lim *et al*. (2) to provide targeted RNA sequencing, which was achieved through an amplicon-based approach using the gene-specific primer of the *Rps29* mRNA and the 3’ adaptor primer. We could clearly detect two peaks representing long poly(A) tails (>40 nt) and short poly(A) tails (10–20 nt) in oocytes before hormonal stimulation (Figure 3A, −1-MA), as observed using Sanger sequencing (Figure 2C, −1-MA). The long poly(A) tails in unstimulated oocytes were not uridylated (Figure 3B, −1-MA, >40), whereas the short poly(A) tails (10–20 nt) were highly uridylated (>60%) (Figure 3B, −1-MA, ≤40). Following hormonal stimulation, these two populations decreased significantly and a new peak (0–10 nt) appeared (Figure 3A, +1-MA). As shown in Figure 2C, the tails of the new peak were uridylated (Figure 3B, +1-MA, ≤40). These results suggest that 1-MA stimulation led to shortening of both long poly(A) tails and short uridylated poly(A) tails, followed by new uridylation of short poly(A) tails during meiosis resumption.

**Figure 3.**
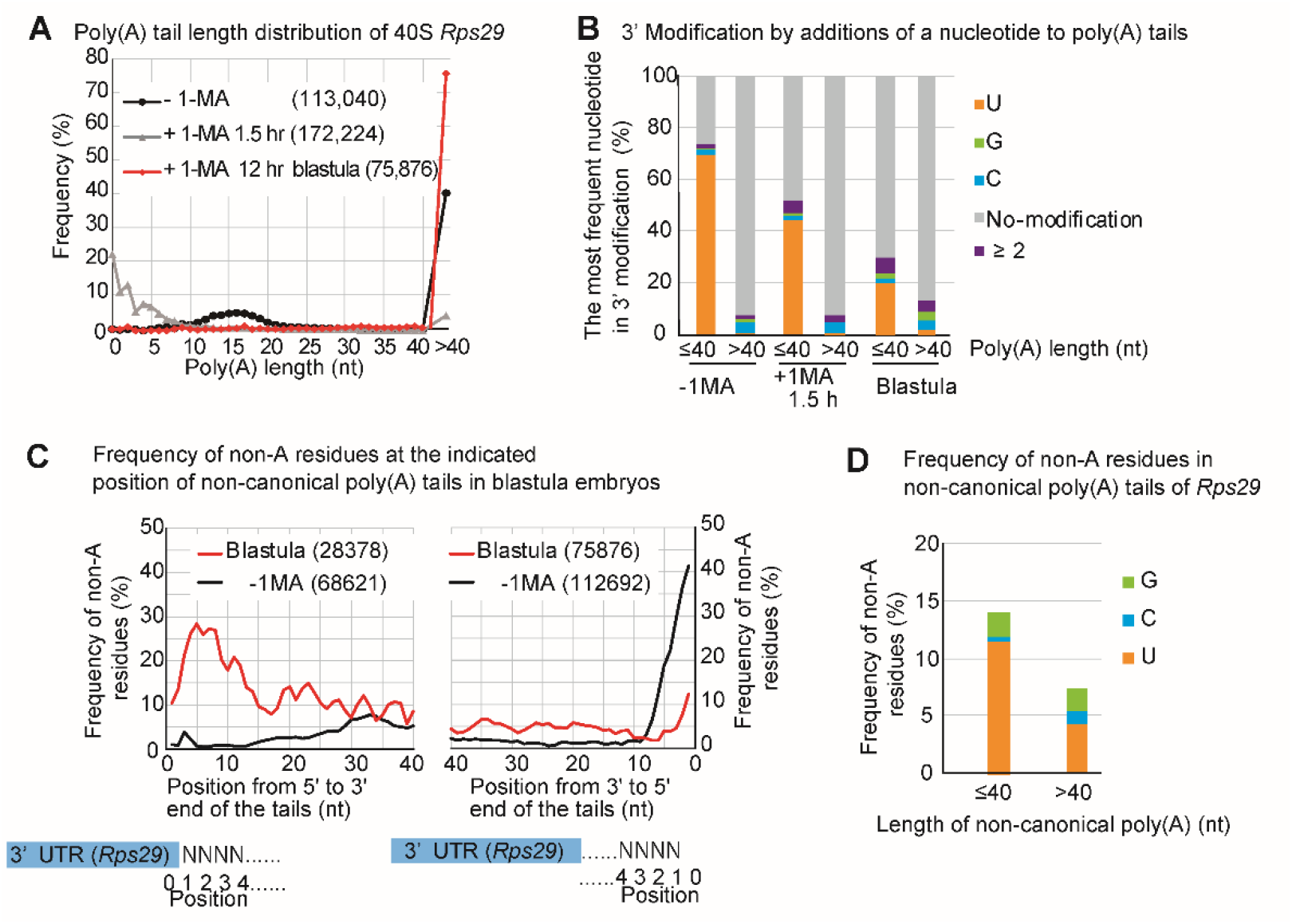
Targeted TAIL-Seq of *Rps29* mRNA. A. Distribution of poly(A) tail lengths of *Rps29* mRNA from oocytes without 1-MA treatment (−1-MA), with 1-MA treatment for 1.5 h (+1-MA 1.5 h), and with 1-MA treatment followed by insemination to obtain blastula embryos at 12 h. Relative frequencies (Y axis, %) were calculated by dividing the number of detected reads that have indicated poly(A) lengths by the total number of reads that have poly(A) tails. Frequencies of poly(A) tail length exceeding 40 nucleotides are plotted at the right side (>40). The number of reads is shown in parentheses. B. Relative frequencies of the most frequent nucleotide in additional modifications at the 3’ end of *Rps29* mRNA. Using each *Rps29* mRNA read, the most frequent nucleotides, such as U, G, and C, were determined. Relative frequencies (Y axis, %) were calculated by dividing the number of reads with the most frequent nucleotides by the total number of reads with the indicated lengths of poly(A) tails. The mRNA that contains tail lengths fewer than or equal to 40 nucleotides (≤) and over 40 (>) are compared in each stage of oocytes and embryos. No modification; neither poly(A) tail nor additional modifications were present at the end of poly(A) tails. ≥2; two or more nucleotides comprised the most frequent nucleotides in the mRNA. C. Distribution of relative frequencies of non-A residues in tails of *Rps29* mRNA from blastula embryos and unstimulated oocytes. At the indicated position of the tails, the relative frequencies of the non-A residues (Y axis, %) were calculated by dividing the number of reads carrying the non-A residues by the total number of reads. The distribution of frequencies of non-A residues is shown at the indicated position in tails from 5’ to 3’ (left panel) and from 3’ to 5’ (right panel). The numbers of reads are shown in parentheses. D. Relative frequencies of non-A residues in the non-canonical poly(A) tails of *Rps29* mRNA. The relative frequencies of the non-A residues (Y axis, %) were calculated by dividing the number of each non-A residue (G, C, and U) in the tails of all reads by the number of tail lengths of all reads that have indicated lengths of poly(A) tails. The mRNA that carries tail lengths of fewer than or equal to 40 nucleotides (≤) and over 40 (>) are compared at each stage of oocytes and blastula embryos.

Because the frequency of non-A residues appeared to be higher in the 5’ region of non-canonical poly (A) tails of *Rps29* mRNA in blastula embryos (Figure 2C, +1-MA, 12 h), we calculated the percentages of non-A residues at the indicated position of residues in tails from 5’ to 3’ using the results of TAIL-Seq (Figure 3C, left panel). As expected, approximately 30% of the residues were non-A in the 5’ region, with the percentages decreasing gradually as the number of residues increased towards the 3’ direction. Furthermore, when we calculated percentages of non-A residues in tails from 3’ to 5’ in blastula embryos (Figure 3C right panel), the frequencies of non-A residues from blastula embryos were lower than those from 5’ to 3’ (Figure 3C, left panel), indicating that the 3’ end region of the non-canonical poly(A) tails contains more A residues than the 5’ end region. The frequencies of uridine or guanine were higher than those of cytosine in the non-canonical poly(A) tails (Figure 3D). These results suggest that non-canonical poly(A) construction was initiated by the formation of a U-rich tail followed by A-rich tail elongation. When we calculated the percentages of non-A residues in unstimulated oocytes from 3’ to 5’ (Figure 3C right panel), the frequency of non-A residues at the 3’ end in oocytes before hormonal stimulation was approximately 40%, which may be due to uridylated short poly(A)s (10–20 nt) carrying additional U tails (Figure 2C, endogenous −1-MA; Figure. 3B, −1-MA, ≤40). Similar results were obtained when a targeted TAIL-Seq was performed using the specific primer for *Rpl27a* mRNA (Supplemental Figure S3).

### Injected 3’ UTR of Rps29 mRNA behaves similarly to endogenous mRNA

To confirm whether 40S *Rps29* mRNA that contains non-canonical poly(A) tails originated from maternal mRNA that contains long canonical poly(A), but not from zygotic mRNA transcribed during embryogenesis, we synthesised the 3’ UTR of *Rps29* mRNA harbouring both a unique sequence tag and a long poly(A) tail and injected this 3’ UTR into oocytes (Figure 4A). If maternal *Rps29* mRNA was deadenylated after 1-MA stimulation and re-adenylated in morula and/or blastula embryos, the exogenous 3’ UTRs of *Rps29* mRNA carrying long poly(A) tails would be deadenylated after hormonal stimulation, followed by re-elongation of the poly(A) tails.

**Figure 4.**
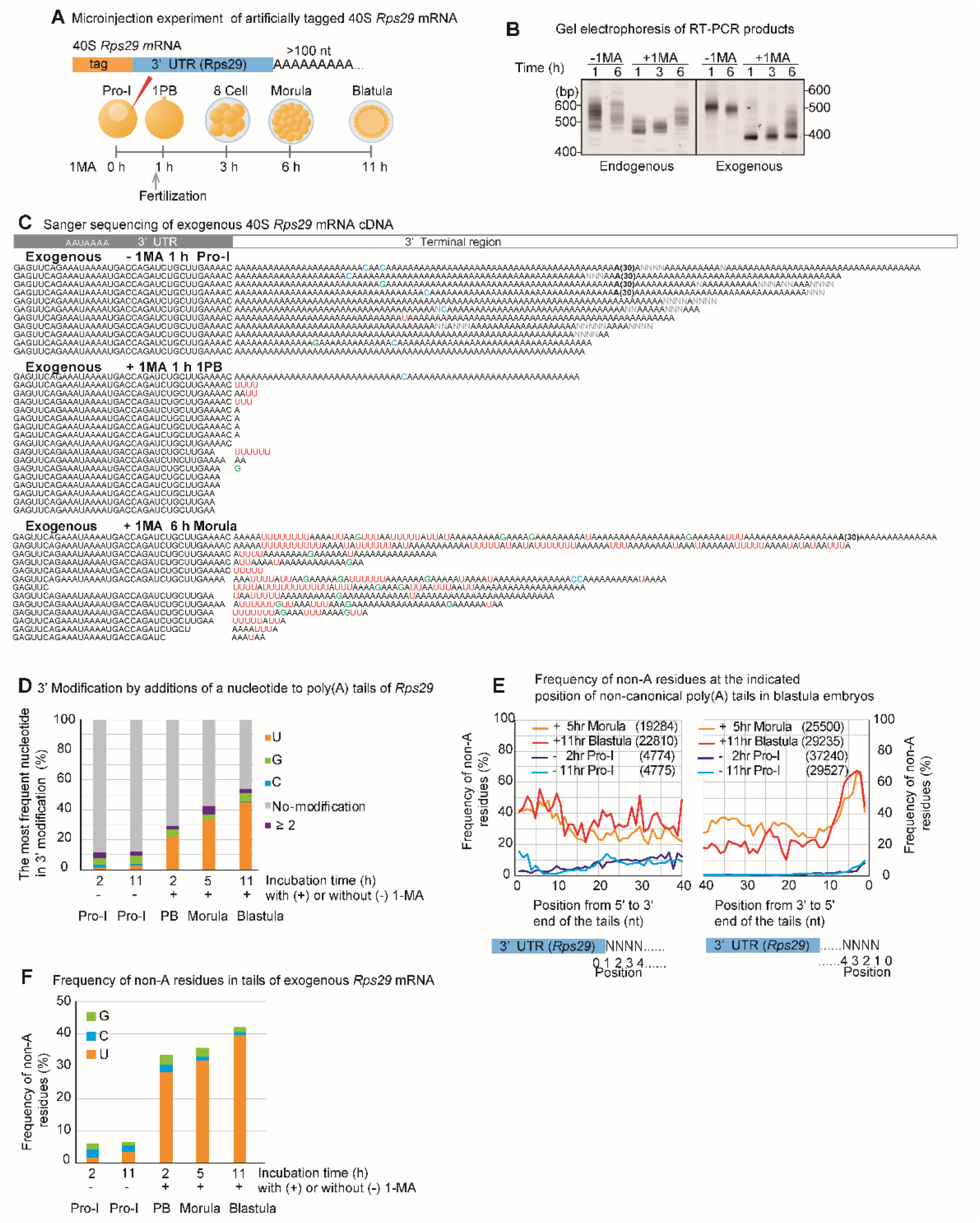
Deadenylation, uridylation, and non-canonical poly(A) elongation of exogenous ribosomal protein *Rps29* mRNA. A. Experimental scheme. Tag-labelled 40S *Rps29* mRNAs carrying long poly(A) tails without modification were synthesised and injected into oocytes without 1-MA treatment (−1-MA). To obtain developing embryos, oocytes stimulated with 1-MA were inseminated before first polar body formation. Total RNAs from oocytes with or without 1-MA stimulation were purified at the indicated times (1, 3, and 6 h), whereas stimulated oocytes at 1 h and unstimulated oocytes were not inseminated. B. Measurement of the tail length of *Rps29* mRNA. Total RNAs purified from oocytes or embryos were subjected to TGIRT template-switching reaction (Supplemental Figure S1, B left panel). RT-PCR was conducted using the 3’ adaptor reverse primer and *Rps29* mRNA -specific forward primer. The PCR products were then subjected to polyacrylamide gel electrophoresis and visualised using SYBR-Green I staining. The left and right panels show changes in the tail lengths of endogenous and exogenous *Rps29* mRNA, respectively. Similar results are obtained using 2 animals. C. Sequencing results of the 3’ terminal region of exogenous 40S *Rps29* mRNA from oocytes at Pro-I without 1-MA treatment [Exogenous −1-MA], stimulated oocytes at 1 h following 1-MA treatment [Exogenous +1-MA 1 h], and morula embryos (6 h after 1-MA treatment) [Exogenous +1-MA 6 h]. ‘(30)’ in the sequences indicates ‘AAA…AAA’ containing 30 nucleotides. D. Relative frequencies of the most frequent nucleotide in additional modifications at the 3’ end of exogenous *Rps29* mRNA. Using each read of exogenous *Rps29* mRNA, the most frequent nucleotides, such as U, G, and C, were determined. Relative frequencies (Y axis, %) were calculated by dividing the number of reads carrying the most frequent nucleotide by the total number of reads from oocytes and embryos at the indicated time following 1-MA stimulation. No modification; neither poly(A) tail nor additional modifications were present at the end of poly(A) tails. ≥2; two or more nucleotides comprised the most frequent nucleotides in the mRNA. E. Distribution of relative frequencies of non-A residues in tails of exogenous *Rps29* mRNA from oocytes and embryos. At the indicated position of the tails, the relative frequencies of the non-A residues (Y axis, %) were calculated by dividing the number of reads carrying the non-A residues by the total number of reads. The distribution of frequencies of non-A residues is shown at the indicated position in tails from 5’ to 3’ (left panel) and from 3’ to 5’ (right panel). The numbers of reads are shown in parentheses. Yellow, morula embryos (5 h after 1-MA treatment) Red, blastula embryos (11 h after 1-MA treatment). Purple, Pro-I oocytes without 1-MA stimulation (2 h after injection) Blue, Pro-I oocytes without 1-MA stimulation (11 h after injection). F. Relative frequencies of non-A residues in canonical and non-canonical poly(A) tails of exogenous *Rps29* mRNA. The relative frequencies of the non-A residues (Y axis, %) were calculated by dividing the number of each non-A residue (G, C, and U) in the tails of all reads by the number of tail lengths of all reads.

As expected, electrophoresis of the RT-PCR products of the endogenous and exogenous *Rps29* mRNA tails showed their lengths decreased similarly at 1 h following hormonal stimulation and increased in the morula embryos at 6 h after hormonal stimulation (Figure 4B). In addition, Sanger sequencing revealed that the long poly(A) tails of exogenous RNA injected into oocytes were not only removed but also uridylated after hormonal stimulation (Figure 4C, +1-MA 1 h). Thus, 1-MA stimulation shortened both endogenous and exogenous poly(A) tails of *Rps29* mRNA, followed by uridylation. Moreover, the exogenous RNA carried U-rich non-canonical poly(A) tails in morula embryos (Figure 4C +1-MA 6 h), as observed for endogenous mRNA (Figure 2C +1-MA 12 h). These results support the hypothesis that the deadenylated and uridylated maternal RNAs following 1-MA stimulation were re-adenylated to generate U-rich non-canonical poly(A) tails in morula embryos. This re-adenylation signal is likely to be included in the 3’ UTR of *Rps29* mRNA because the exogenous mRNA carried only a 3’ UTR sequence and a unique sequence tag.

Targeted TAIL-Seq showed that the frequency of uridine addition at the 3’ ends of the injected *Rps29* mRNA increased following hormonal stimulation (Figure 4D 2–11 h (+) 1-MA), whereas no increase was observed in unstimulated oocytes (Figure 4D 2 h, −1-MA, 11 h, −1-MA). The frequency of non-A residues in the 5’ region was approximately 40% in the exogenous *Rps29* mRNA tails from morula and blastula embryos (Figure 4E left panel: +1-MA 5 h, +1-MA 11 h), which was comparable to that of endogenous *Rps29* mRNA tails (Figure 3C left panel, blastula). However, the 3’ region contained more non-A than endogenous A residues (Figure 4E right panel: +1-MA 5 h, +1-MA 11 h; Figure 3C blastula), likely due to the shorter tails of the injected RNAs (Figure 4B, right panel, +1-MA, 6 h) compared to those of endogenous mRNAs, which may contain U-rich sequences without poly(A) tails. The frequency of uridine was higher than guanine or cytosine in the exogenous non-canonical poly(A) tails (Figure 4F).

### The AAUAAA recognition site of the cleavage and polyadenylation specificity factor (CPSF) is not required for re-polyadenylation

Sanger sequencing revealed that some exogenous *Rps29* mRNAs in morula embryos or endogenous *Rpl27a* mRNAs in blastula embryos did not contain the polyadenylation signal (PAS) ‘AAUAA’ (Figure 4C, morula, lane 7; Supplemental Figure S2 blastula, lanes 4 and 9), suggesting that they were polyadenylated even after trimming of the 3’ UTR including PAS. To confirm that PAS is not required for re-polyadenylation in morula or blastula embryos, we synthesised ΔPAS RNA in which the U of AAUAAA was mutated to G (Figure 5A). When we injected this construct into oocytes, followed by hormonal stimulation and fertilisation, ΔPAS RNA behaved similarly during development to wild-type RNA carrying PAS (Figure 5B and C). The targeted TAIL-Seq of injected ΔPAS mRNA showed that modification of the 3’ terminal region occurred similarly to that in mRNA carrying PAS (Supplemental Figure S4A-C).

**Figure 5.**
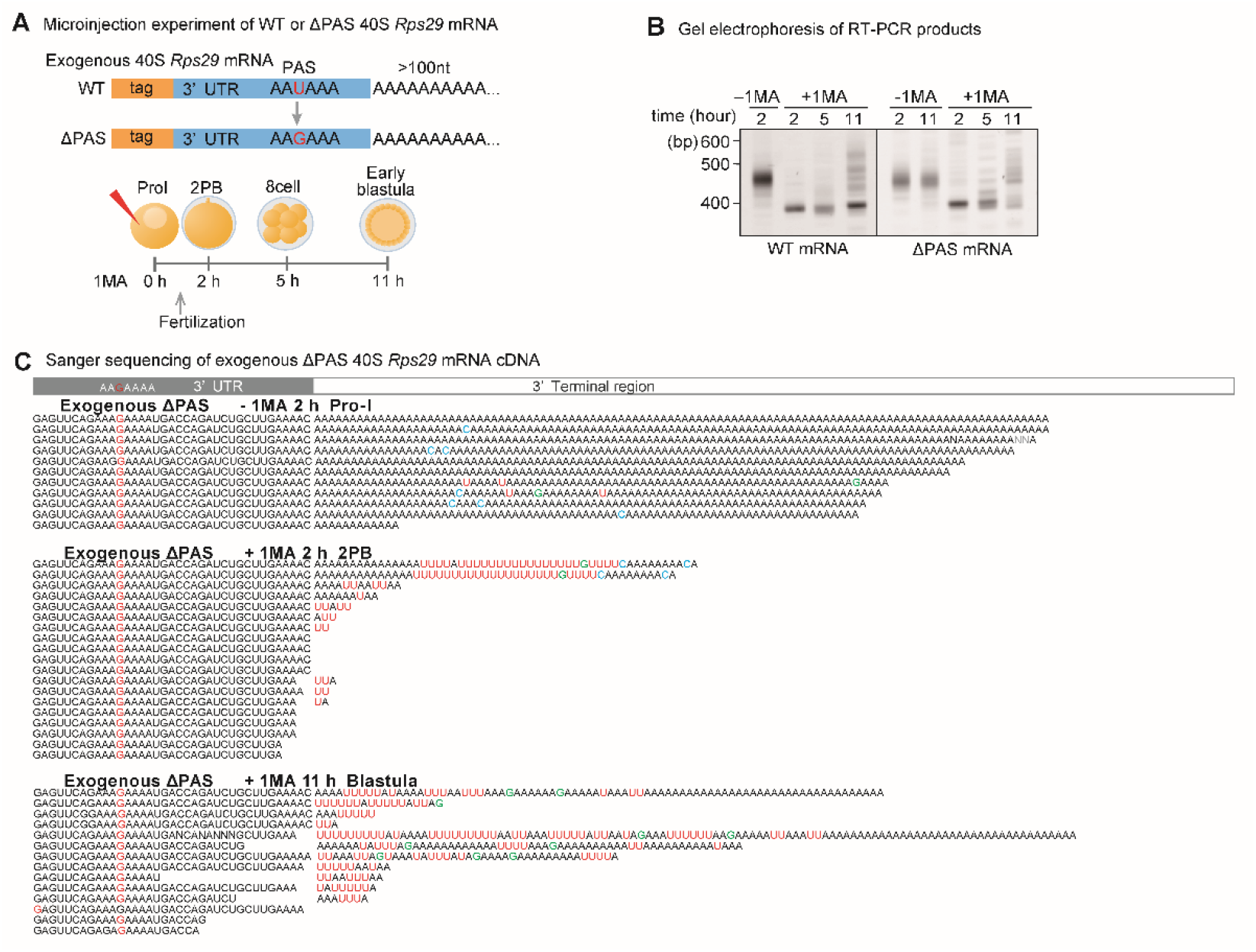
Deadenylation, uridylation, and non-canonical poly(A) elongation of exogenous ribosomal protein Δ PAS *Rps29* mRNA. A. Experimental scheme. Tag-labelled 40S wild-type and Δ PAS *Rps29* mRNAs were synthesised and injected into oocytes without 1-MA treatment (−1-MA). Stimulated oocytes with 1-MA were inseminated before first polar body formation. At the indicated time, the total RNA was purified. B. Measurement of the tail length of *Rps29* mRNA. Total RNA purified from oocytes or embryos was subjected to TGIRT template-switching reaction (Supplemental Figure S1, B left panel). RT-PCR was conducted using the 3’ adaptor reverse primer and Tag-specific forward primer. The PCR products were then subjected to polyacrylamide electrophoresis and visualised using SYBR-Green I staining. The left and right panels show changes in tail lengths of exogenous wild-type and Δ PAS *Rps29* mRNA, respectively. Similar results are obtained using 2 animals. C. Sequencing results of the 3’ terminal region of exogenous Δ PAS *Rps29* mRNA purified from oocytes at Pro-I without 1-MA treatment at 2 h following injection of the mRNA [Exogenous ΔPAS −1-MA 2 h], stimulated oocytes at 2 h following 1-MA treatment [Exogenous +1-MA 2 h], and blastula embryos (11 h after 1-MA treatment) [Exogenous +1-MA 11 h].

### Non-canonical poly(A) tailed mRNA is translationally active

To investigate whether the non-canonical(A) tail enhances the translational activity of mRNAs, we applied a protein expression system for immature starfish oocytes injected with starfish SGK mRNA (22). Because the anti-sfSGK antibody was more sensitive than the anti-FLAG tag antibody, we used sfSGK as a protein tag. We synthesised four types of sfSGK mRNAs: carrying a long poly(A) tail (>100 nt), a short poly(A)tail (4 nt), and carrying each of two types of non-canonical poly(A) tail from blastula *Rps29* cDNA clones encoding nc1 and nc2. (Figure 6A). Western blot analysis of oocytes injected with each mRNA to evaluate translational activity revealed that mRNAs with long poly(A) tails and nc1 tails supported substantial expression of exogenous SGK protein (Figure 6B). These results suggest that the A-rich poly(A) region (31 nt) in the non-canonical poly(A) tail of nc1 mRNA exhibited translational activity comparable to that of the long poly(A) tail (>100 nt). Furthermore, when the reporter mRNA inserted between the 5’ and 3’ UTR of 40S *Rps29* with a canonical poly(A) tail was synthesised and injected into stimulated oocytes, followed by fertilisation (Fig. 6C), the embryos showed an increase in translational activities at the blastula stage (Fig. 6 D). Because the changes in the exogenous mRNA poly(A) tail length in injected embryos mimicked those in endogenous *Rps29* mRNA (Fig. 6 E; Fig. 2 B), these results support the hypothesis that deadenylated and uridylated *Rps29* mRNAs are recycled, re-polyadenylated, and translated to produce ribosomal proteins in blastula-stage embryos.

**Figure 6.**
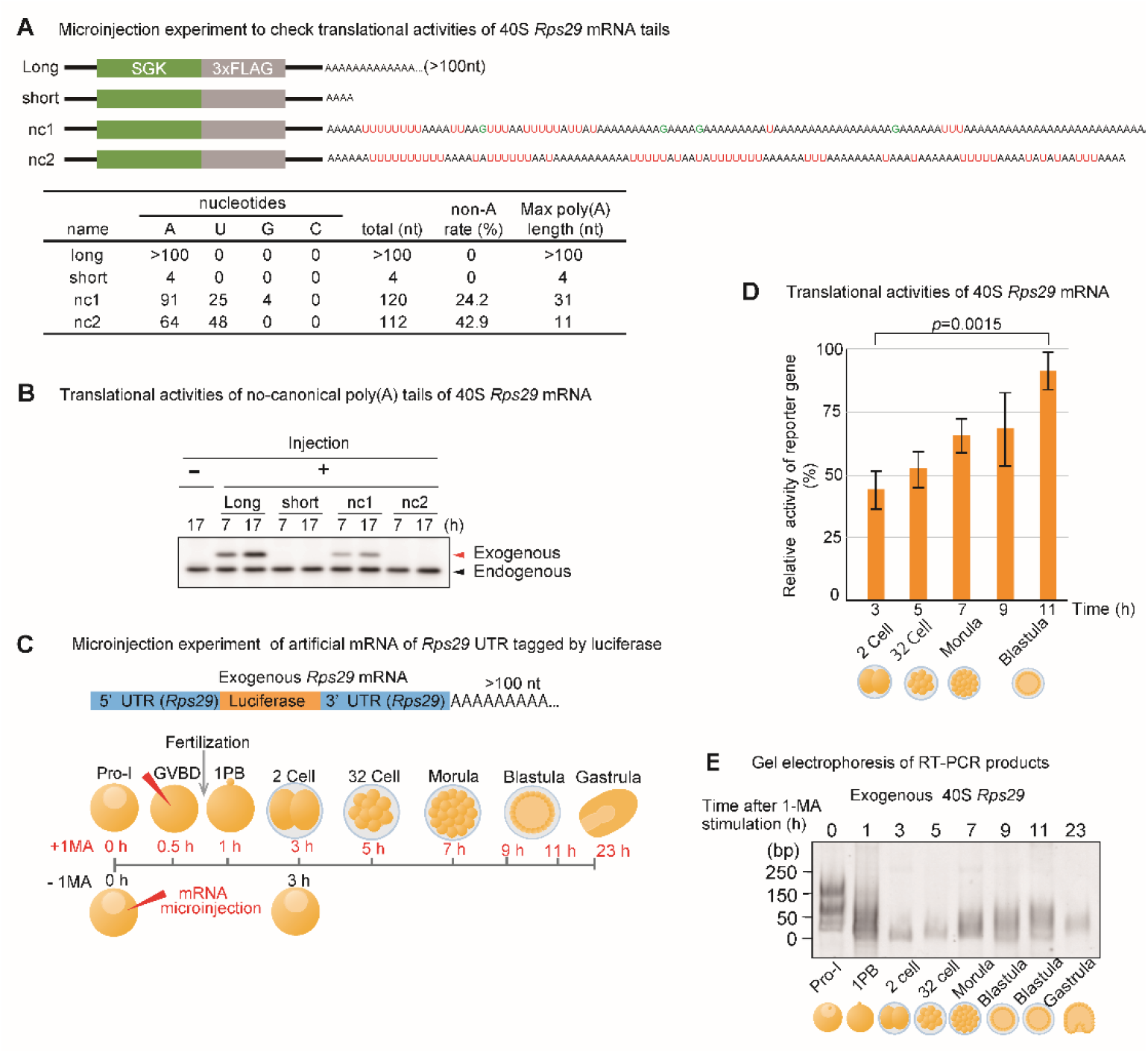
Translational activities of non-canonical poly(A) tails. A. Experimental scheme. Tag(SGK)-labelled 40S *Rps29* mRNA having canonical long poly(A) tails, short poly(A) tails, and non-canonical long and short poly(A) tails were synthesised and injected into oocytes without 1-MA treatment. The table shows the number of A, T, G, and C nucleotides in the tails. B. Western blotting of oocytes injected with mRNA carrying an SGK tag and non-canonical poly(A) tails of 40S *Rps29* mRNA. At the indicated time after injection of exogenous mRNA, oocytes were treated with sample buffer, followed by polyacrylamide gel electrophoresis and western blotting using an anti-SGK antibody. Arrowheads (endogenous) indicate endogenous SGK in oocytes. Similar results are obtained using 3 animals. C. Experimental scheme. Reporter luciferase mRNA between the 5’ and 3’ UTR of 40S *Rps29* with a canonical long poly(A) tail was injected into oocytes with or without 1-MA treatment. D, E. 1-MA-stimulated oocytes were inseminated after GVBD to start embryonic development and used to determine the reporter activities (D) (median ± SEM) (n = 3), and the length of poly(A) tails (E). Translational activity of oocytes, which were injected with the reporter mRNA with a long poly(A), without 1-MA stimulation at 3 h was accepted as 100% and Student’s t-test was used to determine the significance between the results observed after 3 and 11 h (D). Similar results were obtained using two animal models (D and E).

## Discussion

In this study, using invertebrate starfish, we investigated the 3’ -terminal modification of maternal mRNAs for ribosomal proteins and *cyclin B* during oocyte maturation and embryonic development. Starfish *cyclin B* mRNA carrying a long canonical poly(A) tail was deadenylated and uridylated at the blastula stage, followed by decay, indicating that uridylation of *cyclin B* mRNA promotes mRNA degradation in starfish embryos, as previously demonstrated in vertebrate embryos (38) (Figure 7A, upper panel, #5). Conversely, *Rps29* and *Rpl27a* mRNAs producing starfish ribosomal proteins in oocytes before hormonal stimulation did not decay even when they were deadenylated and uridylated following hormonal stimulation or fertilisation (Figure 7A, upper panel, #1). In addition, they were re-polyadenylated, forming translationally active U-rich non-canonical poly(A) tails at the blastula stage (Figure 7A, upper panel, #2). To our knowledge, this is the first report demonstrating the re-adenylation of uridylated mRNAs to achieve secondary translational activity for mRNA recycling. Furthermore, uridylated short poly(A) tails in starfish have two fates: degradation (Figure 7. A #5) or poly(A) elongation (Figure 7, #2 and #3). Thus, uridylation of starfish mRNAs in a stably inactive state functions as a bidirectional signpost, towards the formation of unstable, inactive mRNAs for degradation or stable, eventually active mRNAs for translation (Figure 7, lower panel; and Figure 7B).

**Figure 7.**
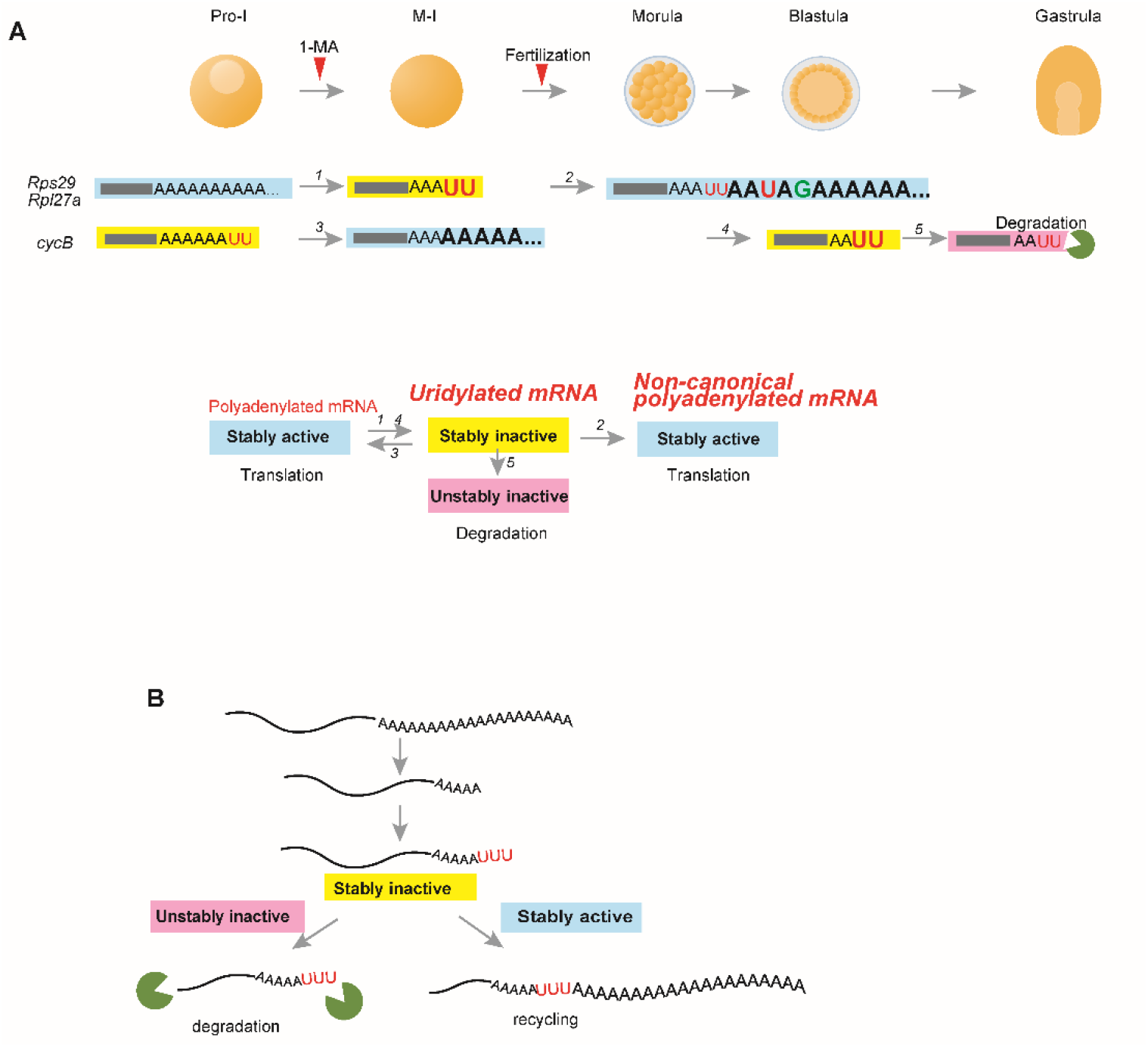
Summary of findings and proposed model. A. Summary of maternal mRNA deadenylation, uridylation, and non-canonical poly(A) elongation in starfish oocytes and embryos. Upper panel: Before hormonal stimulation of 1-MA, oocytes at Pro-I contain maternal ribosomal protein mRNAs (*Rps29*and *Rpl27a*) carrying long poly(A) tails and *cyclin B* mRNAs with uridylated short poly(A) tails. Following the resumption of meiosis of oocytes undergoing nuclear division of germinal vesicle breakdown, the long poly(A) tails of ribosomal protein mRNAs are deadenylated and uridylated (newly added nucleotides are shown in a larger font size). Some uridine residues of *cyclin B* mRNA are trimmed (20), followed by poly(A) elongation. After fertilisation, the uridylated short poly(A) tails of ribosomal proteins in morula embryos are re-elongated, forming non-canonical poly(A) tails. At the blastula stage, *cyclin B* mRNAs are deadenylated and uridylated. They are then degraded before gastrulation. Lower panel: Upper panel summarisation. The arrow numbers correspond to those in the upper panel. Uridylated mRNAs are stable but inactive for translation. Canonical or non-canonical polyadenylation renders inactive mRNAs stably active for translation. Alternatively, uridylated mRNAs become unstable and inactive, followed by decay, as observed in other animals. B. Proposed model for uridylation in starfish. Uridylated mRNAs are required for determining mRNA fate: destruction or recycling.

Although the mechanisms underlying the decision-making processes of degradation or re-polyadenylation of uridylated mRNAs in starfish embryos remain to be fully elucidated, our results indicate that the 3’ UTR is likely involved in the choice of fate and uridylation, because the 3’ UTRs of ribosomal protein mRNAs and *cyclin B* mRNA reproduced the pattern of degradation or re-polyadenylation of endogenous mRNAs at the morula, blastula, and gastrula stages (Figure 1 and 4). Upon resumption of meiosis in *Xenopus*, mouse, and starfish oocytes, cytoplasmic polyadenylation of many maternal mRNAs, including *cyclin B*, is regulated by 3’ UTRs containing the PAS AAUAAA and the CPE, which are bound by CPSF and the CPEB, respectively (16, 39– 41). Subsequently, in *Xenopus* oocytes, the terminal nucleotidyl transferase 2 Gld2, which interacts with CPSF and CPEB, elongates the poly(A) tails (18, 42, 43). Alternatively, mRNAs that do not contain CPE or PAS are deadenylated after hormonal stimulation in both amphibian and mouse oocytes (15, 44, 45). Similarly, maternal mRNAs of starfish ribosomal proteins, *Rps29* and *Rpl27a*, which do not contain CPE, were deadenylated after hormonal stimulation (Figure 2, Supplemental Figure S2). Moreover, re-polyadenylation of the *Rps29* mRNA occurred even in endogenous mRNA trimmed from the 3’ end to the site of PAS, and in exogenous 3’ UTR carrying a mutation in PAS (Supplemental Figure S2 blastula, lanes 4 and 9; Figure 4C, morula, lane 7), suggesting that Gld2 may not be involved in non-canonical poly(A) elongation.

Deadenylated starfish *cyclin B* mRNA was uridylated at the blastula stage, followed by decay, suggesting that terminal uridylyltransferases such as mammalian TUT4/7 (7, 31) may be involved in this uridylation (Figure 7, A #4). In *Xenopus* embryos, degradation of *cyclin B2* mRNA depends on the 3’ UTR (32), which contains a microRNA-427 (miR-427) target sequence, with the zygotic expression of miR-427 inducing *cyclin B2* mRNA deadenylation (46). Similarly, the decay of starfish *cyclin B* mRNA may also be induced by microRNA, because duplex formation of morpholino oligonucleotides mimicking the interaction of microRNA with the mRNA 3’ UTR terminus degrades the mRNA in starfish (23).

In *Xenopus*, many maternal mRNAs are not decayed immediately following deadenylation, but are stable until the blastula stage, several hours after fertilisation (9– 13). Although it remains to be determined how *Xenopus* maternal mRNAs with short poly(A) tails are stabilised after deadenylation, the possibility exists that uridylation of *Xenopus* maternal mRNAs, which do not necessarily cause decay, may occur, similar to the results observed in starfish. Consistent with this, when we analysed the TAIL-Seq data of *Xenopus* embryos kindly provided by Hyeshik Chang and Narry Kim (31), approximately 20% of *Xenopus* 40S *rps29* mRNAs were modified, to be uridylated at stage 5 (Supplemental Figure S5). Thus, some *Xenopus* mRNAs may be stable even when uridylated, suggesting the presence of a stable inactive state of uridylated mRNA in both vertebrates and invertebrates.

Non-A residues are observed more frequently near the 5’ end of poly(A) tails in mouse GV oocytes, as shown using poly(A) inclusive RNA isoform sequencing (PAIso−Seq) (47). Similarly, approximately 30% of the non-A residues were dominantly distributed in the 5’ region of the non-canonical poly(A) tails of the mRNA of the ribosomal protein in starfish embryos (Figure 2C and Figure 3C), suggesting that both animals may utilise similar modification systems. Although the enzymes involved in the 5’ end modifications in mouse oocytes remain to be determined, mammalian TENT4A/4B (48) can mediate mixed tailing of adenylation and guanylation to stabilise mRNAs. Thus, the starfish homologue of TENT4 may generate the 5’ modifications in the non-canonical poly(A) tails. In addition, TENT5 families mediate the cytoplasmic polyadenylation of collagen mRNAs required for bone mineralisation (49) and immunoglobulin mRNAs (50) in mammals. These families may also participate in the non-canonical polyadenylation, exhibiting a high percentage of A residues, especially at the 3’ region in the starfish cytoplasm.

The La-related protein 1 (LARP1), which recognises the UUU-3’ -OH of the terminal motif of transcripts synthesised by RNA polymerase III in nuclei (51), can control the translation and stability of mammalian ribosomal protein mRNAs in the cytoplasm (52, 53). The potential therefore exists that the stability of uridylated ribosomal protein mRNAs in the cytoplasm of early embryos in starfish may also be regulated by molecules such as LARP1.

In this study, we proposed there is a stable inactive state of uridylated mRNAs in starfish. However, it remains to be confirmed whether stably inactive mRNAs with uridylated tails exist in other organisms. Although we could not address this question directly, besides the suggestive results from *Xenopus*, the presence of uridylated mRNA in various cells (6, 30) suggests that additional waiting phases are required to recruit molecule to degrade or possibly re-elongate mRNA tails. Therefore, more studies are needed regarding the recycling of mRNAs (54) or re-elongation of uridylated mRNAs in various animals to address these issues.

## Data availability

The MiSeq sequencing data generated in this study were deposited under BioProject PRJDB9545. The accession numbers of the mRNAs of *Rps29* (40S ribosomal protein) and *Rpl27a* (60S ribosomal protein) are LC535328 and LC535329, respectively.

## Funding

This work was supported by JSPS KAKENHI [Grant Number JP17K07405, 21K06185 to K.C.], the Takeda Science Foundation; and the Cooperative Program provided by the Atmosphere and Ocean Research Institute, University of Tokyo.

## Acknowledgement

We would like to thank Dr Hyeshik Chang and Dr Narry Kim for providing us TAIL-Seq data of *Xenopus* embryos. We would like to thank Editage (www.editage.com) for English language editing.

## Conflict of interest

The authors declare that they have no competing interests.

## Notes

### Competing Interest Statement

The authors have declared no competing interest.

