## Supplemental Tables and Figures for "Recycling of uridylated mRNAs in starfish embryos"

**Table S1.** Primers for RT-PCR

| Figure | Target | Primer name | Sequence (5′ to 3′) |
|---|---|---|---|
| Figure 1 | starfish *cyclin B* | sfcycB_F | GAAAGCCCATCTGATCAAGC |
| | | TGIRT_R | CAGACGTGTGCTCTTCCGAT |
| Figure 2 | starfish *Rps29* | sfRps29_F1 | TCACATCTGCCCCAAAGCAT |
| | | PCR-R&RT | GTCTCTAGCCTGCAGGA |
| Figure 4 | starfish *Rps29* | sfRps29_F2 | TGTGTGGCAATCGTCATGGA |
| | | TGIRT_R | CAGACGTGTGCTCTTCCGAT |
| Figure 4 | exogenous mRNA | exo_tag_F | AAAGATCTGCAGCGTGCGTCA |
| | | TGIRT_R | CAGACGTGTGCTCTTCCGAT |
| Figure 6 | Starfish Rps 29 5’ UTR | 40S 5’ RACE R | TCGGTTGGTGTTGCTGCGCT |
| Figure S2 | *sfRpl27a* | sfRpl27a_F | AAGCCAAGTTCTTCAGCCGA |
| | | PCR-R&RT | GTCTCTAGCCTGCAGGA |

**Table S2** Primers for PCR to construct expression vectors and qPCR

| oligo DNA | sequence (3' to 5') |
|---|---|
| psfcycB_vector_F | GTTCCATCCTACGAATTCCT |
| psfcycB_vector_R | TTTAACTTTTATTTACAAAGACTGACA |
| psfcycB_A25_insert_F | AAATAAAAGTTAAATGGAAATAAAAAAAAAAAAAAAAAAAAAAAAAAAACGCATGTTCCATCCTACGAA |
| psfcycB_A25_insert_R | TTCGTAGGATGGAACATGC |
| psfRps29_WT_vector_F | TGGCGGCCGCTCTAGAAAGATCTGCAGCGTGCGTCAAACCTACTCTAGAAGCAGCTGG |
| psfRps29_WT_vector_R | GTTTTCAAGCAGATCTGGTCATT |
| psfRps29_WT_insert_template | GACCAGATCTGCTTGAAAACAAAAAAAAAAAAAAAAAAAAGAGACGCTAGAACTAGTGGATCCCCC |
| psfRps29_WT_insert_F | GACCAGATCTGCTTGAAAAC |
| psfRps29_WT_insert_R | ATCCACTAGTTCTAGCGTCTC |
| psfSGK_vector_F | TGCAGGTCGACTCTAGAGGA |
| psfSGK_vector_R | AGAAACTTTCTTTTTATTAGGAGCAGATAC |

| | |
|---|---|
| psfSGK_long_insert_F | AAAAAGAAAGTTTCTTCACATTCTAAAAAAAAAAAAAAAAAAAAAAAAAAGTCTTCTGCAGGTCGACTCTA |
| psfSGK_long_insert_R | TAGAGTCGACCTGCAGAAGAC |
| psfSGK_short_insert_F | GTTTCTTCACATTCTAAAAAAAGTCTTC |
| psfSGK_short_insert_R | TAGAGTCGACCTGCAGAAGACTTTTTTT |
| psfSGK_nc1_insert_F | GTTTCTTCACATTCTAAAAATTTTTTTTAAAATTAAGTTTAATTTTTATTATAAAAAAAAAGAAAAGAAAAAAAAATAAAAAAAAAAAAAAAAAGAAAAAATTTAAAAAAAAAAAAAAAAAAAAAAAAAAAAAAAGTCTTCTGCAGGTCGACTCTA |
| psfSGK_nc1_insert_R | TAGAGTCGACCTGCAGAAGAC |
| psfSGK_nc2_insert_F | GTTTCTTCACATTCTAAAAAATTTTTTTTTAAAATATTTTTTAATAAAAAAAAAAATTTTTATAATATTTTTTTAAAAAATTTAAAAAAAATAAATAAAAAATTTTTAAAATATATAATTTAAAAGTCTTCTGCAGGTCGACTCTA |
| psfSGK_nc2_insert_R | TAGAGTCGACCTGCAGAAGAC |
| 18S F qPCR | GCGGCCGAAACGTTTACTTT |
| 18S R qPCR | TTCCATGCTCTGCTGTCCAG |
| Cyclin B F qPCR | CTCAACAACGCACCCAAGT |
| Cyclin B R qPCR | TGAACTTGCTGCTGGCATAC |
| 40S ribosomal protein S29 F qPCR | GGCAATCGTCATGGACTGAT |
| 40S ribosomal protein S29 R qPCR | GTGAACTGCACCCTGAGAAA |
| sfRps29-GST_1F | CGCGGTGGCGGCCGCTCTAGGACCAAATTGTAAAATTTCTTTCTCT |
| sfRps29-GST_1R | TAACCTAGTATAGGGGACATATCCATCTTTTGAAATCCAATGTC |
| sfRps29-GST_2F | ATGTCCCCTATACTAGGTTATTGGA |
| sfRps29-GST_2R | AGCTGCTTCTAGAGTAGGTTTCATTTTGGAGGATGGTCGCC |
| BsmBI-pBlue3' | ATCCACTAGTTCTAGCGTCTC |
| sfRps29-GST_3F | AACCTACTCTAGAAGCAGCTGG |
| sfRps29-GST_3R | CTAGAGCGGCCGCCACC |
| Nluc_FW2 | CTCGTGGACGCCATCATGGTCTTCACACTCGAAG |
| Nluc_RV2 | CTTCTAGAGTAGGTTTTACGCCAGAATGCGTTCG |
| vec_FW2 | AACCTACTCTAGAAGCAGC |
| vec_RV2 | GATGGCGTCCACGAGGTGC |

**Table S3.** Statistics of targeted TAIL-Seq reads.

| Sample description | Target gene | Endogenous/Exogenous | SRA accession | # Raw read pairs | # Valid read pairs |
|---|---|---|---|---|---|
| *Rps29* -1-MA | *Rps29* (40S) | Endogenous | DRR218044 | 159,138 | 113,040 |
| *Rps29* +1-MA 1.5 h | *Rps29* (40S) | Endogenous | DRR218045 | 212,874 | 172,224 |
| *Rps29* +1-MA 12 h Blastula | *Rps29* (40S) | Endogenous | DRR218046 | 124,150 | 75,876 |
| *Rpl27a* −1-MA | *Rpl27a* (60S) | Endogenous | DRR218047 | 112,496 | 52,351 |
| *Rpl27a* +1-MA 1.5 h | *Rpl27a* (60S) | Endogenous | DRR218048 | 117,322 | 64,800 |
| *Rpl27a* +1-MA 12 h Blastula | *Rpl27a* (60S) | Endogenous | DRR218049 | 101,332 | 30,792 |
| WT *Rps29* −1-MA 2 h Pro-I | *Rps29* (40S) | Exogenous (injected) | DRR218050 | 65,670 | 38,789 |
| WT *Rps29* −1-MA 11 h Pro-I | *Rps29* (40S) | Exogenous (injected) | DRR218051 | 52,396 | 30,867 |
| WT *Rps29* +1-MA 2 h PB | *Rps29* (40S) | Exogenous (injected) | DRR218052 | 76,771 | 58,632 |
| WT *Rps29* +1-MA 5 h Morula | *Rps29* (40S) | Exogenous (injected) | DRR218053 | 49,119 | 33,505 |
| WT *Rps29* +1-MA 11 h Blastula | *Rps29* (40S) | Exogenous (injected) | DRR218054 | 54,821 | 32,352 |
| ΔPAS *Rps29* −1-MA 2 h Pro-I | *Rps29* (40S) | Exogenous (injected) | DRR218055 | 96,099 | 59,555 |
| ΔPAS *Rps29* −1-MA 11 h Pro-I | *Rps29* (40S) | Exogenous (injected) | DRR218056 | 68,731 | 42,567 |

| | | | | | |
|---|---|---|---|---|---|
| ΔPAS *Rps29* +1-MA 2 h PB | *Rps29* (40S) | Exogenous (injected) | DRR218057 | 77,342 | 52,094 |
| ΔPAS *Rps29* +1-MA 5 h Morula | *Rps29* (40S) | Exogenous (injected) | DRR218058 | 88,182 | 64,476 |
| ΔPAS *Rps29* +1-MA 11 h Blastula | *Rps29* (40S) | Exogenous (injected) | DRR218059 | 58,726 | 32,017 |
| *cyclin B* TGIRT | *cyclin B* | Endogenous | DRR218060 | 171,337 | 148,143 |
| *cyclin B* T4 RNA ligase | *cyclin B* | Endogenous | DRR218061 | 309,649 | 283,824 |

**A** Schematic model of the anaylsis of targeted TAIL-seq.

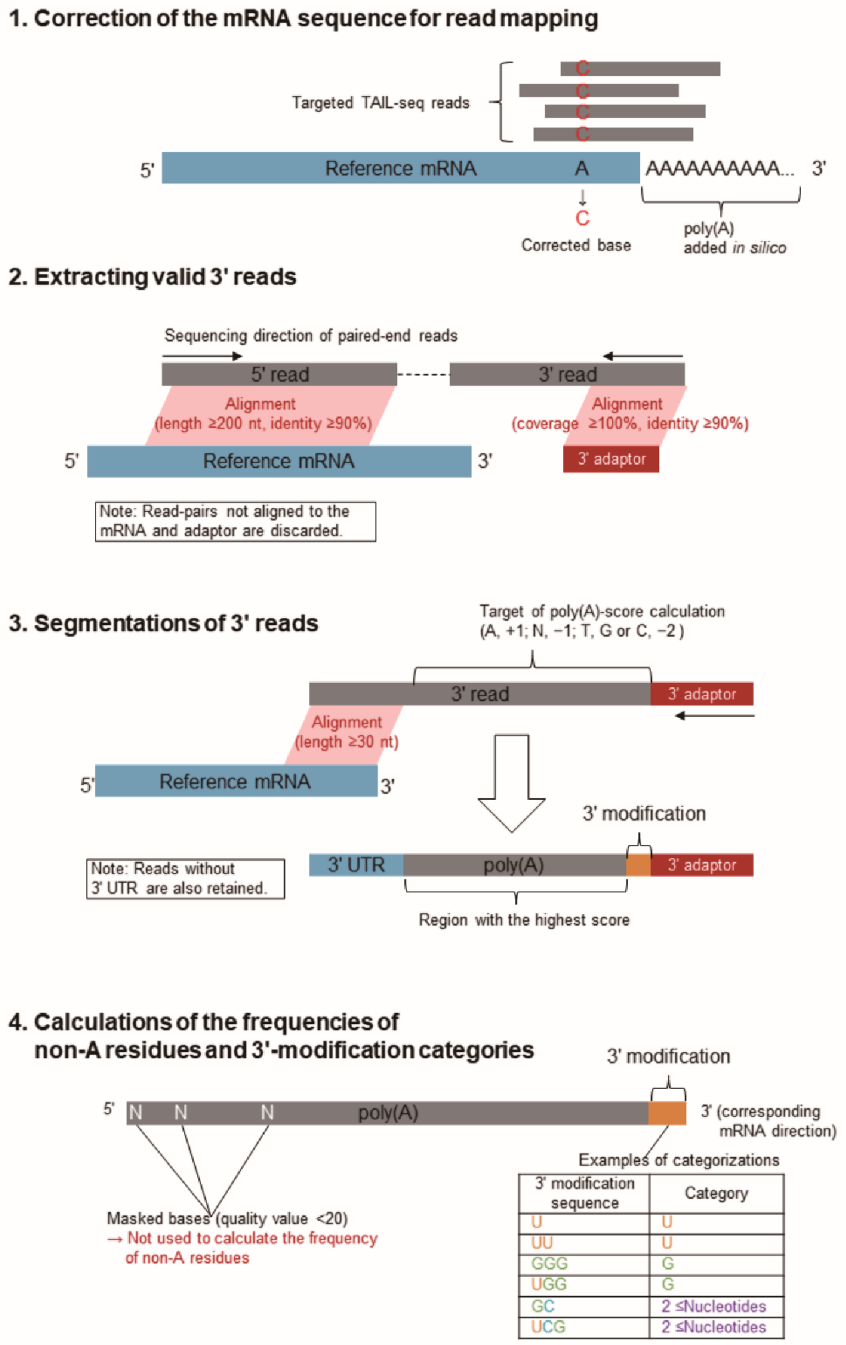

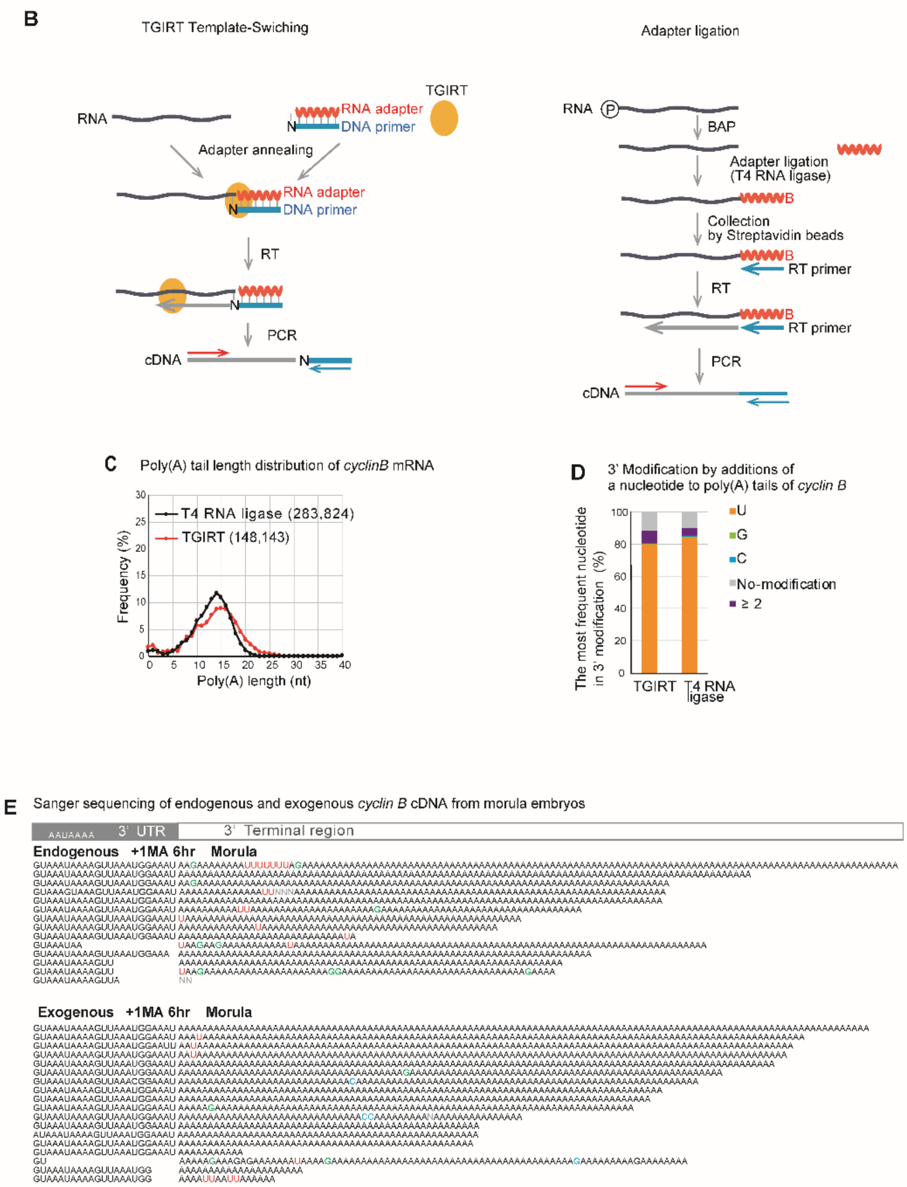


**Supplemental Figure S1.** Experimental schemes, TAIL-Seq and Sanger sequences.

A. Schematic model of the analysis of targeted TAIL-Seq. See detail in the text of Materials and Methods.

B. Experimental scheme of the TGIRT template-switching reaction (left panel) and adaptor ligation (right panel). Left panel: The DNA primer for TGIRT has a single nucleotide 3′ overhang with an equimolar mixture of A, T, G, and C, which can base pair with the 3′ end nucleotide of the target RNA. Reverse transcription with this enzyme, which is a TGIRT template-switching reaction, forms cDNA (55). Right panel: Biotinylated 3′ adaptor was ligated by T4 RNA ligase to the 3′ end of the mRNA, followed by reverse transcription to produce cDNA (20). PCR was conducted using a gene-specific primer and 3′-adaptor primer. These two methods yielded identical results, as shown in B and C (below). The template-switching reaction by TGIRT had good efficiency, requiring 30 oocytes (A, left panel), whereas the method of ligating adapters using T4 RNA ligase (A, right panel) had poor reaction efficiency, requiring 100 or more oocytes. C. Distribution of poly(A) tail lengths of *cyclin B* mRNA in oocytes without 1-MA treatment. The results obtained by the TGIRT template-switching reaction (red) were identical to those obtained by the T4 RNA ligase method (black), when targeted TAIL-Seq of *cyclin B* mRNA was conducted. Relative frequencies (Y axis, %) were calculated by dividing the number of detected reads carrying the indicated poly(A) lengths by the total number of reads containing poly(A) tails. The numbers of reads are shown in parentheses. D. Relative frequencies of the most frequent nucleotide in additional modifications at the 3′ end of *cyclin B* mRNA poly(A) tails. Using each *cyclin B* mRNA read, the most frequent nucleotides, U, G, and C, were determined. Relative frequencies (Y axis, %) were calculated by dividing the number of reads with the most frequent nucleotides by the total number of reads. No modification; neither poly(A) tail nor additional modifications were present at the end of poly(A) tails. ≥2; two or more nucleotides comprised the most frequent nucleotides in the mRNA. E. Sequencing results of the 3′ terminal region of endogenous and exogenous *cyclin B* mRNA from morula embryos (6 h after 1-MA treatment).

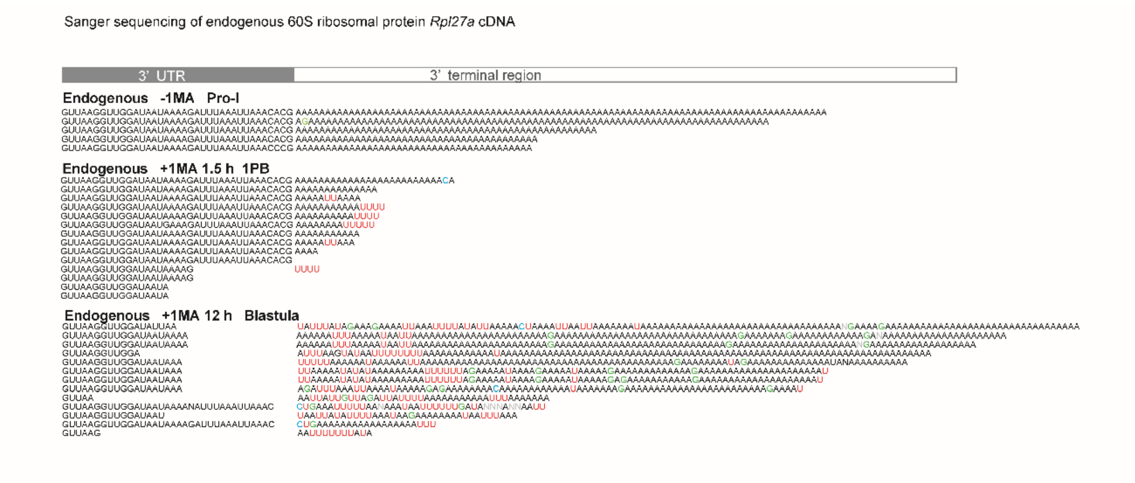


**Supplemental Figure S2.** Sequencing results of the 3′ terminal region of endogenous maternal *Rps27a* mRNA from oocytes at Pro-I without 1-MA treatment [Endogenous −1-MA], stimulated oocytes following first polar body formation (1.5 h after 1-MA treatment) [Endogenous +1-MA 1.5 h], and blastula embryos (12 h after 1-MA treatment) [Endogenous +1-MA 12 h].

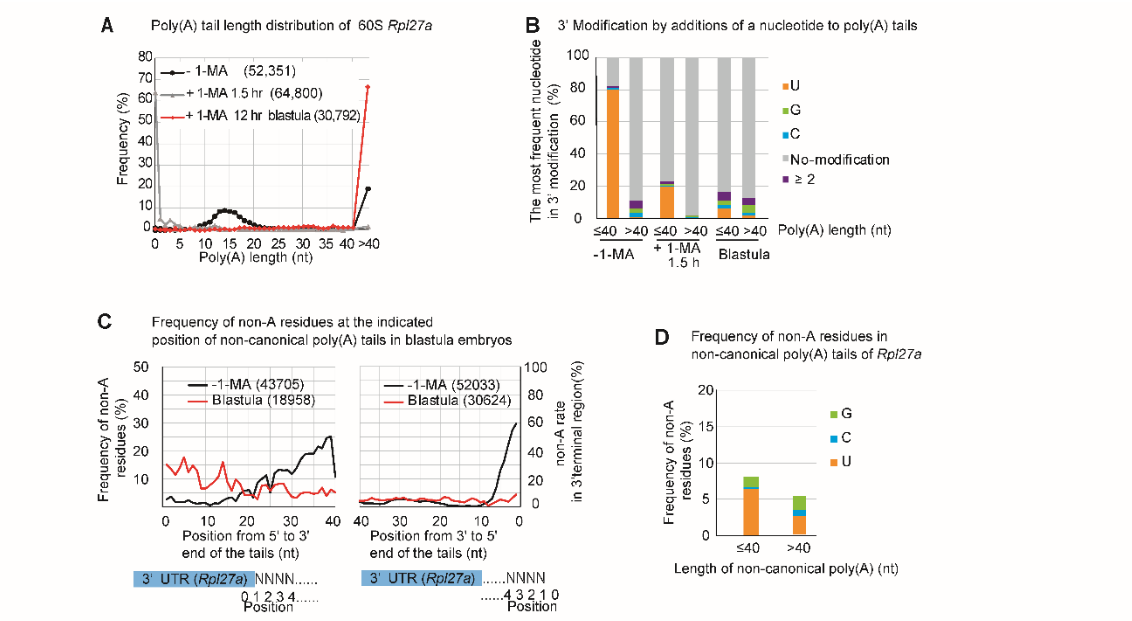


**Supplemental Figure S3**. Targeted TAIL-Seq of *Rpl27a* mRNA.

A. Distribution of poly(A) tail lengths of *Rpl27a* mRNA from oocytes with (+1-MA) or without 1-MA treatment (−1-MA), and from blastula embryos at 12 h following 1-MA treatment. Relative frequencies (Y axis, %) were calculated by dividing the number of detected reads carrying the indicated poly(A) lengths by the total number of reads containing poly(A) tails. Frequencies of poly(A) tail lengths of over 40 nucleotides are plotted on the right side (>40). The numbers of reads are shown in parentheses.

B. Relative frequencies of the most frequent nucleotide in additional modifications at the 3′ end of *Rpl27a* mRNA. Using each *Rpl27a* mRNA read, the most frequent nucleotides, such as U, G, and C, were determined. Relative frequencies (Y axis, %) were calculated by dividing the number of reads with the most frequent nucleotides by the total number of reads carrying the indicated lengths of poly(A) tails. The mRNAs carrying tail lengths of less than or equal to 40 nucleotides (≤), and over 40 (>) were compared in each stage of oocytes and embryos. No modification; neither poly(A) tail nor additional modifications were present at the end of poly(A) tails. ≥2; two or more nucleotides comprised the most frequent nucleotides in the mRNA.

C. Distribution of relative frequencies of non-A residues in the tails of *Rpl27a* mRNA from blastula embryos and unstimulated oocytes. At the indicated position of the tails, the relative frequencies of the non-A residues (Y axis, %) were calculated by dividing the number of reads carrying the non-A residues by the total number of reads. The distribution of frequencies of non-A residues is shown at the indicated position in tails from 5′ to 3′ (left panel) and from 3′ to 5′ (right panel). The numbers of reads are shown in parentheses.

D. Relative frequencies of non-A residues in the non-canonical poly(A) tails of *Rpl27a* mRNA. The relative frequencies of the non-A residues (Y axis, %) were calculated by dividing the number of each non-A residue (G, C, and U) in the tails of all reads by the number of tail lengths of all reads that have indicated lengths of poly(A) tails. The mRNAs carrying tail lengths of less than or equal to 40 nucleotides (≤), and over 40 (>) were compared in each stage of oocytes and embryos.

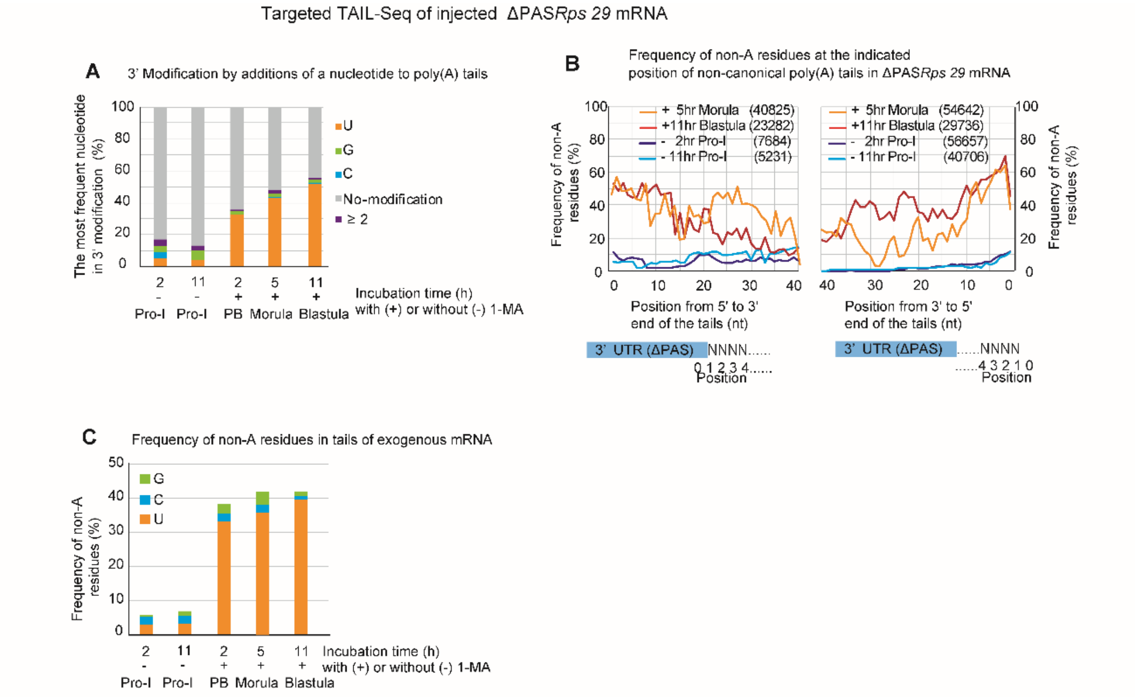


**Supplemental Figure S4** Targeted TAIL-Seq of Delta PAS *Rps29* mRNA. A. Relative frequencies of the most frequent nucleotide in additional modifications at the 3′ end of the exogenous Δ PAS *Rps29* mRNA. Using each exogenous Δ PAS *Rps29* mRNA read, the most frequent nucleotides, such as U, G, and C, were determined. Relative frequencies (Y axis, %) were calculated by dividing the number of reads carrying the most frequent nucleotide by the total number of reads from oocytes and embryos at the indicated time following 1-MA stimulation. No modification, either tail or additional modifications, was present in the mRNA. ≥2; over two nucleotides were the most frequent nucleotides in the mRNA. B. Distribution of relative frequencies of non-A residues in tails of exogenous Δ PAS *Rps29* mRNA from oocytes and embryos. At the indicated position of the tails, the relative frequencies of the non-A residues (Y axis, %) were calculated by dividing the number of reads carrying the non-A residues by the total number of reads. The distribution of frequencies of non-A residues is shown at the indicated position in tails from 5′ to 3′ (left panel) and from 3′ to 5′ (right panel). The numbers of reads are shown in parentheses. Yellow, morula embryos (5 h after 1-MA treatment) Red, blastula embryos (11 h after 1-MA treatment). Purple, Pro-I oocytes without 1-MA stimulation (2 h after injection) Blue, Pro-I oocytes without 1-MA stimulation (11 h after injection). C. Relative frequencies of non-A residues in canonical and non-canonical poly(A) tails of exogenous Δ PAS *Rps29* mRNA. The relative frequencies of the non-A residues (Y axis, %) were calculated by dividing the number of each non-A residue (G, C, and U) in the tails of all reads by the number of tail lengths of all reads.

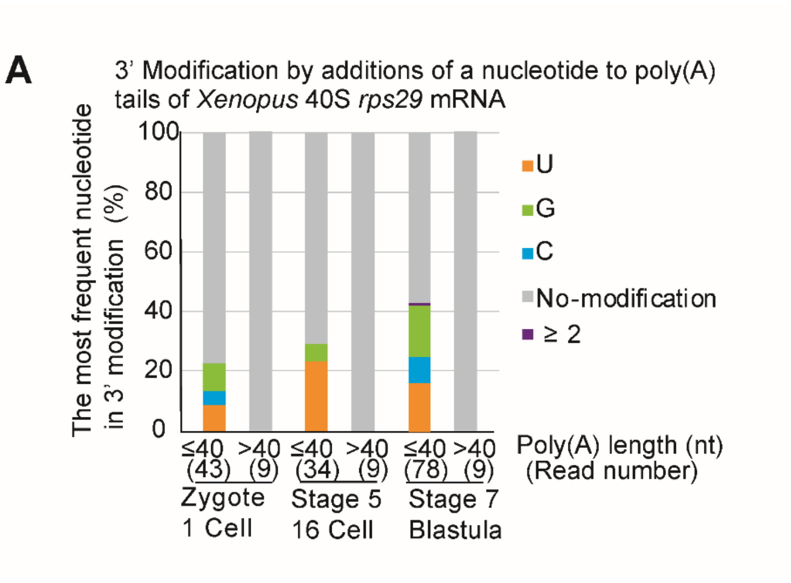


**Supplemental Figure S5.** Relative frequencies of the most frequent nucleotide in additional modifications at the 3′ end of *Xenopus Rps29* mRNA. Using TAIL-Seq of *Xenopus* embryos (31), the most frequent nucleotides, such as U, G, and C, were determined. Relative frequencies (Y axis, %) were calculated by dividing the number of reads that carry the most frequent nucleotide by the total number of reads from oocytes and embryos at the indicated stage. No modification; neither poly(A) tail nor additional modifications were present at the end of poly(A) tails. ≥2; two or more nucleotides comprised the most frequent nucleotides in the mRNA.
